# MDA5 generates compact ribonucleoprotein complexes via ATP-dependent single-stranded RNA loop extrusion

**DOI:** 10.1101/2024.08.27.609867

**Authors:** Salina Quack, Sourav Maity, Pim P. B. America, Misha Klein, Alba Herrero del Valle, Rahul Singh, Quinte Smitskamp, Flavia S. Papini, Chase P. Broedersz, Wouter H. Roos, Yorgo Modis, David Dulin

## Abstract

Long double-stranded (ds) RNA in the cytosol acts as a potent inflammatory molecule recognized by the receptor MDA5, triggering the innate immune response. Mutations affecting MDA5 ATPase activity lead to severe pathological conditions. MDA5 nucleoprotein filament assembly-disassembly dynamic is proposed to regulate dsRNA recognition, though the exact mechanism remains unclear. Here, we employed magnetic tweezers to monitor the assembly and manipulate MDA5 filaments at the single dsRNA level. Following a slow nucleation event, MDA5 assembles cooperatively and directionally into (partial) filaments and utilizes ATP hydrolysis to compact dsRNA via single-stranded (ss) RNA loop extrusion, even against a significant opposing force. This compacted state is further stabilized by oligomerization of MDA5’s caspase recruitment domain (CARD) and requires high force to be disrupted. ssRNA gaps impaired compaction, suggesting a new mechanism for dsRNA recognition. We propose that MDA5 mediated dsRNA compaction captures viral dsRNA, preventing further usage for viral replication.

## Introduction

Long double-stranded (ds) RNA is a product of RNA and DNA virus replication ^1,2^, and its presence in the cytosol triggers an innate immune response ^3^. Moreover, it has recently been shown that dsRNA may also originate from cellular dysfunctions, and triggers the same innate immune response ^4,5^. Recognition of dsRNA in the cytosol is achieved by pattern-recognition receptors, such as the melanoma differentiation-associated gene-5 (MDA5) ^6^. MDA5 is one of the three RIG-I (retinoic acid-inducible gene I)-like receptors that also includes RIG-I, which binds to di- or tri-phosphorylated 5’-ends, and LGP2, which preferentially binds to blunt-ended dsRNA ^6^. MDA5 proofreads RNA in the cytosol to detect and signal the presence of virus- or retroelement-derived long dsRNA (>100 bp), but generally not structured cellular RNAs ^7^.

MDA5 contains a central RIG-I-like superfamily 2 RNA helicase module flanked by two N-terminal caspase activation and recruitment domains (2CARD) and a C-terminal domain (CTD) ^8,9^. The CTD and helicase module associate with the phosphodiester backbone of dsRNA in a ring-like conformation to cooperatively assemble into helical filaments on long (> 500 bp) dsRNA ^8–16^. It has been proposed that filament assembly plays a central role in recognition and proofreading, through a competition between cooperative assembly and ATP-hydrolysis dependent dissociation, enabling the selection of dsRNAs longer than 500 bp ^11,12,17^. In addition, structural studies have shown that conformational changes coupled to ATP hydrolysis, including an increase in MDA5 footprint on dsRNA from 14 to 15 bp, confer a mechanical proofreading activity, thereby promoting dissociation from imperfectly base-paired RNAs ^8,9^ (**Fig. 1A**). The increase in filament footprint from consecutive rounds of ATP hydrolysis by adjacent MDA5s can displace tightly bound proteins from dsRNA ^18,19^. Furthermore, ATP hydrolysis also enables MDA5 monomer long range translocation on dsRNA ^20^. Following the ATP hydrolysis step, the 2CARDs from the MDA5 filament oligomerize ^12,14^ and the resulting oligomer thereafter nucleates the helical fibril formation at the mitochondrial outer membrane through association with the CARD domain of the mitochondrial antiviral signaling (MAVS) proteins ^21,22^. The MAVS CARD oligomers subsequently activate antiviral interferon responses ^16,21,23^, which also results in a positive feedback loop that increases MDA5 expression and its intra-cytosolic concentration ^24–26^.

**Fig. 1:**
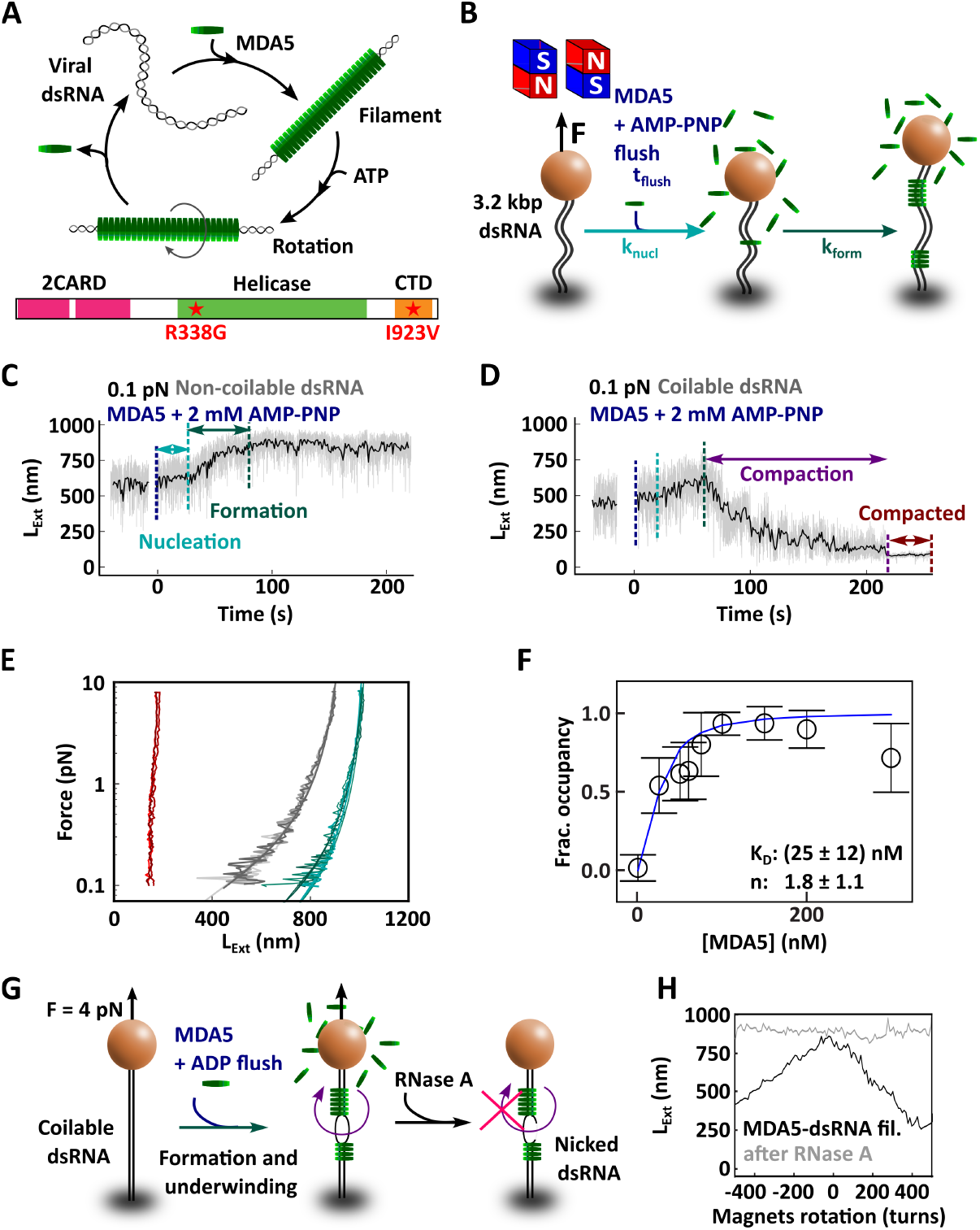
MDA5 assembles cooperatively on long dsRNA in the presence of either ADP or AMP-PNP to form stable and stiff filaments. **(A)** Schematic of the current model of MDA5-RNA activation and of the different domains composing MDA5. MDA5 (green disk) recognizes viral dsRNA (grey) and cooperatively assembles into filaments. Upon ATP hydrolysis, MDA5 monomers rotate and can subsequently dissociate from the dsRNA. Dynamic assembly and disassembly select for long dsRNA. **(B)** Schematic of the magnetic tweezers assay to monitor MDA5 filament formation. **(C, D)** Time trace of MDA5 filament formation with 100 nM MDA5 and 2 mM AMP-PNP on either **(C)** a non-coilable 3.2 kbp dsRNA or **(D)** a coilable 3.2 kbp dsRNA tether, respectively, as illustrated in (B). The vertical dashed lines indicate the end of each phase: flushing of MDA5 with AMP-PNP (blue), nucleation (teal) and filament formation (green). The raw (58 Hz) and time averaged (1 Hz) traces are represented in grey and black, respectively. **(E)** Three repetitions of the force-extension experiment on either a dsRNA tether (grey), a MDA5 filament in the presence of AMP-PNP on non-coilable dsRNA (green), or a coilable dsRNA (red), respectively. The solid lines are worm-like chain model fits to the data (**Equation 2, Materials and Methods**). **(F)** Fractional occupancy (black circles) of MDA5-dsRNA filament according to **Equation 6** fitted with a Hill model (blue line) to obtain the K_*n*_ and the Hill coefficient n (**Equation 1**). The error bars are one standard deviation from 1000 bootstrap samples. **(G)** Description of the experiment demonstrating dsRNA underwinding upon MDA5 binding. The MDA5 filament was assembled on a coilable tether, at 4 pN force, in the presence of ADP and with the magnets clamping the magnetic bead in rotation. Excess of MDA5 was then flushed out and RNase A injected in the flow cell. Rotation-extension experiment was subsequently performed to determine tether coilability. **(H)** Rotation extension curve of a coilable MDA5-dsRNA filament before (black) and after RNase A treatment (grey) as described in (G).

Mutations in the gene encoding MDA5 that compromise its proofreading ability have been linked to interferonopathies ^27,28^, including Aicardi-Goutières syndrome ^29–31^, and Singleton-Merten syndrome ^32^. These pathogenic mutations often alter the ATPase activity of MDA5’s helicase domain, indicating its central role in dsRNA recognition and proofreading. The MDA5 ATPase activity is dependent on dsRNA binding and is tightly regulated for efficient and accurate dsRNA recognition. Indeed, the gain-of-function mutant R338G MDA5 (mouse MDA5, R337G in human) has only 5% of the ATPase activity of WT MDA5. The reduced dissociation rate of MDA5 from dsRNA leads to increased stability of MDA5-dsRNA signaling complexes ^31^. Conversely, the loss-of-function mutant I923V ^11,33^ is ATPase hyperactive and forms less stable filaments. Although the initial steps in RNA recognition by MDA5 have been studied in detail, key questions remain regarding how MDA5 filaments bound to dsRNA assemble into active signaling complexes. In particular, how MDA5 employs ATP hydrolysis to discriminate long dsRNAs from shorter ones remains unanswered. To understand how MDA5 forms filament and utilizes ATP hydrolysis to process dsRNA, high resolution and real-time data monitoring of this process on single long dsRNA molecules are needed.

Here, we employed high-throughput single-molecule magnetic tweezers and high-speed AFM to investigate how MDA5 assembles and processes dsRNA. We confirm that, in absence of ATP, MDA5 filament formation on dsRNA is cooperative and is rate-limited by an initial slow nucleation step. Furthermore, in absence of ATP, we show that MDA5 underwinds dsRNA upon binding. MDA5 forms a partial filament and utilizes ATP hydrolysis to unwind and compact the dsRNA tether by extruding ssRNA loops, even against opposing forces as high as 4 pN. The loop-extruded MDA5-RNA complex is further stabilized and compacted through CARD-CARD interactions and requires 20 pN force to break apart, providing a new pathway to capture viral dsRNA. This study provides an additional mechanism for how MDA5 neutralizes long dsRNA and prevent further usage for viral replication.

## Results

### A single-molecule magnetic tweezers assay to monitor MDA5 filament formation on long dsRNA

We established a high-throughput magnetic tweezers assay to investigate the assembly and mechanochemical properties of MDA5 filaments on long dsRNA ^34–36^. Here, magnetic beads were tethered to the bottom coverslip of a flow chamber via a ∼3.2 kbp long dsRNA (**Fig. 1B**) ^37^. The tether is flanked by two handles, i.e. one with multiple biotin molecules to attach to the streptavidin-coated magnetic bead and the other one with multiple digoxygenin molecules to bind to the anti-digoxigenin coated glass surface of the flow chamber (**Materials and Methods**) ^37^. Several attachment points at both ends rendered fully double-stranded tethers coilable ^38,39^ (**Materials and Methods**). A pair of permanent magnets located above the flow chamber applied an attractive force and constrained the rotation of the magnetic bead ^34,36,37^ (**Materials and Methods**). Hundreds of magnetic beads were imaged simultaneously on a large chip CMOS camera via a home-built inverted microscope. The three dimensional position of the beads was estimated in real-time with nanometer resolution using a custom LabVIEW interface ^40^. The extension of the tether as a function of force was measured by continuously varying the distance between the magnets and the magnetic bead ^41^. These experiments are coined ‘force-extension experiments’ throughout the manuscript and were performed on either bare dsRNA or MDA5 filaments. Around 40% of the dsRNA tethers monitored in the flow chamber were coilable and were used to perform torque-dependent investigations of either bare dsRNA or MDA5-filaments during rotation-extension experiments. The torque applied to the magnetic beads was used to transfer any amount of positive or negative turns into the RNA while measuring the extension of the tether ^38,42–44^.

### MDA5 assembles directionally and cooperatively into stiff and stable filaments in absence of ATP

We investigate MDA5 filament formation and its mechanical properties under force using either a non-hydrolysable form of ATP (AMP-PNP) (**Fig. 1CD**) or the product of ATP hydrolysis (ADP) (**Fig. S1A**). Before adding MDA5 to the flow chamber, the coilability of the dsRNA tethers was determined by performing a rotation-extension experiments at a force of ∼4 pN. At such a force, a coilable dsRNA tether is either constant or decreases in extension when adding either negative or positive turns, respectively (**Fig. S1B**) ^42^. Thereafter, the force was adjusted to 0.1 pN to maximize the change in tether extension upon filament formation, as the filament is expected to be stiffer than bare dsRNA. At this force, bare dsRNA is only stretched at ∼50% of its contour length (*L*_*C*_ = 900 *nm*), and conformational fluctuations of the tether are sufficiently large to occasionally bring distant sections of the filament in contact. Following MDA5 injection into the flow chamber and a subsequent nucleation time, both non-coilable and coilable tethers show an initial increase in extension, indicating the formation of a MDA5 filament. This was followed by either a stable extension of the former (**Fig. 1C**) or a rapid decrease in extension of the latter (**Fig. 1D**). Focusing on the non-coilable tethers first (**Fig. 1C**), we extracted the MDA5 filament nucleation rate and the filament formation rate as a function of MDA5 concentration (**Fig. S1C** and **Fig. S1D**, respectively). If MDA5 binding to the RNA was limiting the nucleation reaction – assuming MDA5 concentration in the flow cell remains constant during the experiment –, we would have expected a monotonic increase in the nucleation rate with MDA5 concentration. However, upon varying MDA5 concentration from 25 to 300 nM, we found that the nucleation rate increased from (0010 ± 0.001) s^-1^ to (0.062 ± 0.010) s^-1^ (mean rate ± standard deviation) and saturated above 150 nM (**Fig. S1C, Table S1**). As the fraction of traces that nucleated during the flush of the proteins was constant above 150 nM MDA5 (**Fig. S1E**), we concluded that the nucleation rate was limited by an event occurring after MDA5 binding to dsRNA, such as accommodating dsRNA into MDA5. We now define the filament formation rate as the average of the local slope of the tether extension obtained from a sliding window (**Material and Methods**). The MDA5 filament formation rate increases exponentially with concentration, i.e. from (0.7-- ± 0.1) monomers/s at 25 nM MDA5 to (8 ± 1) monomer/s (mean rate ± standard deviation, assuming a 14 bp footprint per MDA5 monomer ^8,9^) at 300 nM MDA5 (**Fig. S1D, Table S1**). The filament formation rate is significantly larger than the nucleation rate, supporting a cooperative MDA5 filament formation ^45^, in agreement with previous reports ^11,12,15^.

We performed dynamic force-extension measurements to characterize the mechanical properties of the filament under force, first on the bare dsRNA tether preceding MDA5 injection into the flow chamber, and on the same tether after the MDA5-filament’s extension levelled off (**Fig. 1E, Fig. S1F-N**) ^41^. Fitting the inextensible worm-like chain model ^46^ to the dynamic force-extension traces (**Materials and Methods**), we extract a contour length *L*_*C*_ and a persistence length *L*_*p*_ for every tether (**Fig. S1OP, Table S2**). The dsRNA force-extension data are consistent with previous measurements performed in a buffer containing magnesium, i.e. *L*_*C*_ ≈ 900 *nm* and *L*_*p*_ ≈ 45 *nm* (**Fig. S1OP, Table S2**) ^42,47^. The filaments exhibit a contour length similar to dsRNA (**Fig. 1E, Fig. S1O, Table S2**), while the persistence length increases by ∼4-fold, i.e. up to ∼200 nm (**Fig. S1P, Table S2**). To test the stability of MDA5 filaments in the absence of ATP hydrolysis, we flushed out excess MDA5 proteins with a reaction buffer containing 2 mM of either AMP-PNP or ADP. We subsequently performed force-extension experiments every 30 min during the two hours experiment (**Fig. S2AB**). As the filament’s force-extension curve is largely unchanged over time, we conclude that MDA5 forms stable filaments in standard reaction buffer with either AMP-PNP or ADP (**Fig. S2AB, Table S3**). There is no difference in experiments performed with either AMP-PNP or ADP in those experiments and the following experiments were therefore performed with ADP. In absence of adenosine, MDA5 can form filaments that disassemble over time. This is evidenced by force-extension traces taken at later time points aligning with the bare dsRNA force-extension trace (**Fig. S2C, Table S3**). Adding 0.7 M NaCl in the reaction buffer also resulted in filament disassembly (**Fig. S2D, Table S3**). Furthermore, the MDA5-dsRNA filaments can withstand forces up to ∼50 pN in reaction buffer containing ADP (**Fig. S2E**). Overall, the filaments assembled with either AMP-PNP or ADP are stable in the presence of physiological salt concentration.

To further elaborate on MDA5 binding affinity to dsRNA and cooperative assembly into filaments, we fit a Hill model to the fractional occupancy α extracted from the increase in persistence length of the tether as a function of MDA5 concentration ^48^ (**Materials and Methods**):

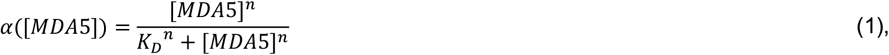

with *K*_*D*_ being the apparent dissociation constant, [*MDA*5] being the concentration of MDA5, and *n* being the Hill coefficient to the fractional occupancy. For wild-type MDA5, we obtain *K*_*D*_ = (25 ± 12) *nM* and *n* = (1.8 ± 1.1) (**Fig. 1F**), confirming a cooperative assembly of MDA5 filament (*n* > 1) ^11,12,15,17^. Increasing the force to 1 pN lowers the noise in the traces and pauses become apparent in the filament formation traces (**Fig. S2F**). If a filament can grow only in one direction, it would grow until reaching an obstacle, e.g. another filament extremity or an attachment point. Further growth would then only be possible if another (slow) nucleation event occurred, resulting in a pause in the tether extension increase. We therefore interpret the pause as a unidirectional filament formation (**Fig. S2F**), in line with previous reports ^17^.

### MDA5 underwinds dsRNA upon binding in absence of ATP hydrolysis

We investigated the torsional properties of the MDA5-dsRNA filament by performing rotation-extension experiments at different forces in the presence of ADP (**Fig. S3A-D**). The MDA5 filament rotation-extension is strongly asymmetric for forces up to 2 pN (**Fig. S3A**), i.e. a rapid decrease in extension upon positive supercoiling, but a slow decrease in extension upon negative supercoiling (**Fig. S3B**). Above 4 pN, the rotation-extension curves are symmetric. When adding negative turns, the extension decreases by 1 nm per added turn and is largely force independent. In contrast, when adding positive turns, the extension decrease per added turn goes from (15 ± 2) *nm/turn* to (0.74 ± 0.06) *nm/turn* when increasing the force from 0.1 to 8 pN, respectively (**Fig. S3C, Table S4**). We note that non-coilable dsRNA tethers are rendered coilable upon MDA5 filament assembly, suggesting that MDA5 monomers can bridge over the nick(s) in the dsRNA backbone (**Fig. S3D**).

Following nucleation on a coilable dsRNA tether, the extension slowly increases (**Fig. 1D, Fig. S3EF, Table S5**) and before reaching the same height as with non-coilable tethers, it starts compacting at a rate of ∼5 nm/s (**Fig. S3F**). All the tethers eventually reach a minimum extension, leading to a tight and irreversibly compacted form recognizable by its significant reduction in the z-axis fluctuation of the bead height (**Fig. S3GH**) and the force-extension measurement (**Fig. 1E**).

Given the difference with non-coilable tethers (**Fig. 1C, Fig. S1A**), we hypothesize that the compaction phase observed with coilable tethers originates from dsRNA being either overwound or underwound upon MDA5 binding. In the former case, the tether would present no single-stranded (ss) RNA region and would therefore be insensitive to RNase A (**Fig. S3IJ**). In the latter case, the dsRNA region of the tether that are not covered by MDA5 filament would melt at forces above 1 pN ^42^ (**Fig. S1B, Fig. S3I**), rendering them sensitive to cleavage by RNase A (**Fig. S3K**). As MDA5 protects dsRNA against RNase degradation ^49^, only free dsRNA that is unwound when negatively supercoiled, would be accessible to RNase A. To this end, we performed rotation-extension experiments on MDA5-dsRNA filaments before and after flushing RNase A into the flow chamber, while keeping the force at 4 pN throughout the experiment (**Fig. 1G**). The addition of RNase A in the flow chamber resulted in the loss of coilability of the tethers (**Fig. 1H**). We therefore conclude that MDA5 underwinds dsRNA in the absence of ATP hydrolysis.

### ATP-hydrolysis induces MDA5 filament compaction stabilized by CARD-CARD interactions

To evaluate how ATP hydrolysis impacts the structural organization of a filament, we pre-assembled the filament in the presence of ADP and subsequently introduced a buffer containing ATP into the flow chamber (**Fig. 2AB**). We immediately noticed a decrease in extension of the tethers after injecting ATP (**Fig. 2B**). We coined this phase ‘compaction’ due to its signature reduction in noise level. The traces also showed abrupt increase in the trace – coined ‘rupture’ events – during the compaction phase (at ∼200 s and ∼300 s in **Fig. 2B**). The compacted filament formed in presence of ATP are very stable and can withstand forces up to (20 ± 4) pN (mean ± standard deviation) (**Fig. S4A**).

**Fig. 2:**
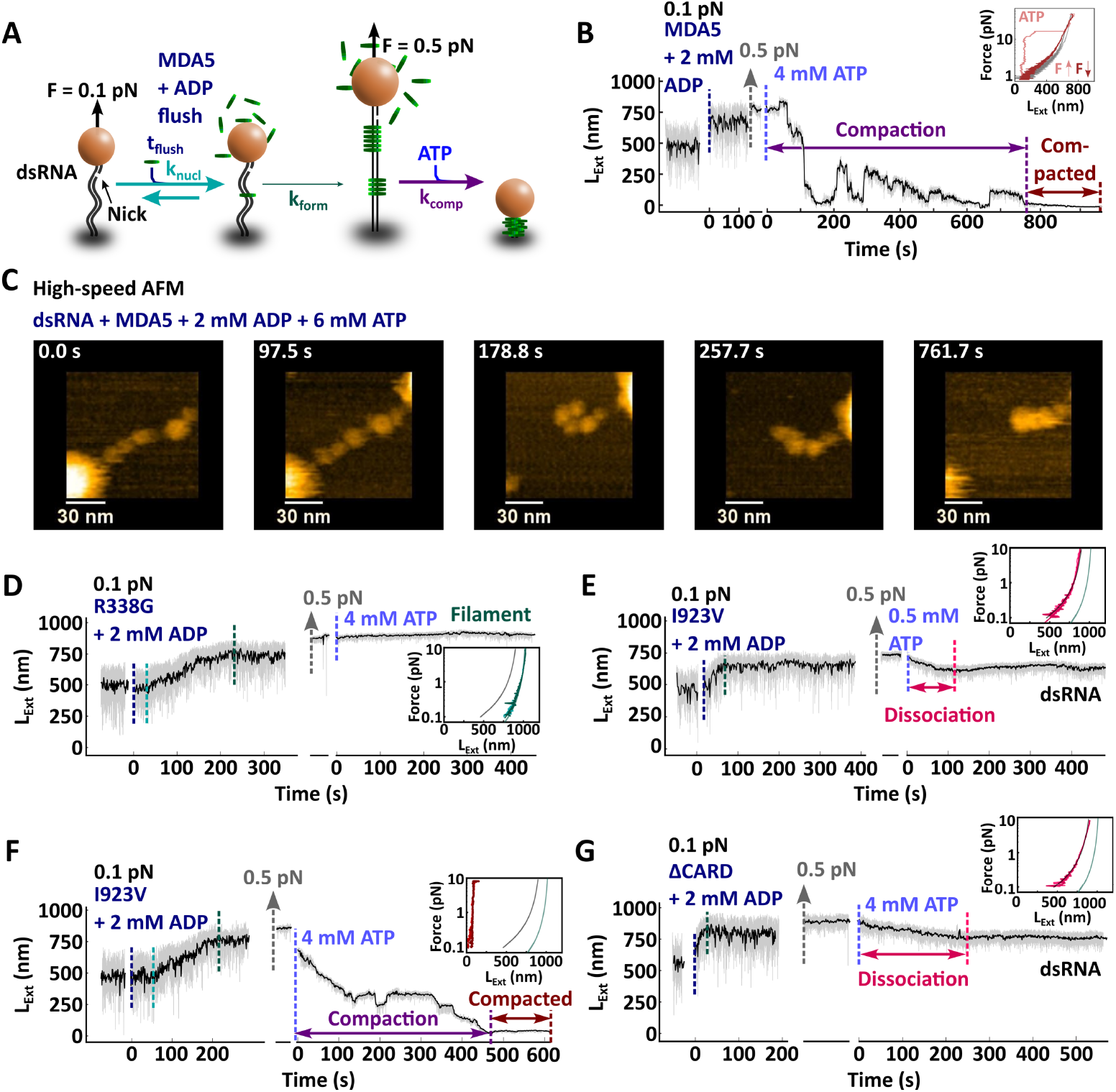
MDA5 filament compaction is ATP-hydrolysis dependent and CARD-CARD interactions are essential to tighten the MDA5-RNA complex. **(A)** Description of the experiment monitoring the filament formation upon injection of 100 nM MDA5 with 2 mM ADP at 0.1 pN force, followed by a force increase to 0.5 pN and a reaction buffer containing 4 mM ATP addition, which removed the MDA5 proteins in excess. **(B)** Time trace of the filament formation and compaction for the experiment described in (A). The vertical dashed lines indicate the end of each phase: flushing (dark blue), nucleation (teal), filament formation (green), filament compaction (purple) and compacted filament (dark red). The raw (58 Hz) and time averaged (1 Hz) traces are represented in grey and black, respectively. Inset: force-extension of a bare dsRNA tether (grey), and a compacted MDA5 filament when increasing (light red) and subsequently decreasing force (dark red). **(C)** Snapshots of HS-AFM video (**Video S1**) of a complex of dsRNA and 50 nM MDA5 after addition of 6 mM ATP (time reset to 0 s). The dsRNA-MDA5 complex was obtained in the presence of 2 mM ADP (**Fig. S4FG**). Imaging rate 300 milliseconds per frame. **(D)** Time trace of the filament formation with MDA5 R338G as described in (A, B). **(E, F)** Time trace of the filament formation with MDA5 I923V as described in (A, B), with addition of either **(E)** 0.5 mM ATP or **(F)** 4 mM ATP. **(G)** Time trace of the filament formation with MDA5 ΔCARD as described in (A, B). Inset in (D, E, F, G): force-extension traces of MDA5 filament after addition of ATP, resulting in either no compaction (green), full compaction (red) or dissociation (pink). Non-extensible WLC fits to force-extension experiments for dsRNA (grey line) and MDA5 filament (teal line). Statistics and WLC fits parameters values are provided in **Table S7**.

We notice that the compaction rate increases exponentially with increasing concentration of ATP (**Fig. S4B, Table S6**) and decreases with force (**Fig. S4C, Table S6**). Furthermore, we observe many rupture events at decreasing force and increasing ATP concentration (**Fig. S4DE, Table S6**). However, such events do not occur once the filament is compacted (**Fig. S4D**). We propose that MDA5 protomers located far from each other within an MDA5 filament can come in close contact during the compaction phase due to thermal motion at forces lower than 1 pN. These fluctuations enable intermolecular CARD-CARD interactions, with occasional rupture events indicating that established bounds broke, but eventually leading to a stable compacted state. Consistent with this model, at higher forces, where the tether extension is close to its contour length, we see a decrease in the number of rupture events per trace (**Fig. S4E**). At higher ATP concentrations we observe more rupture events per trace (**Fig. S4E**), indicating that ATPase activity of MDA5 promotes long distance intermolecular CARD-CARD interactions. In conclusion, MDA5 employs ATP hydrolysis to compact the filament – even against a significant force – into a tight ribonucleoprotein complex further stabilized by CARD-CARD interactions. In absence of force, long distance CARD-CARD interactions facilitate the compaction process.

We then deployed high-speed atomic force microscopy (HS-AFM) ^50,51^ to visualize MDA5’s interactions with dsRNA upon ATP hydrolysis. We pre-assembled MDA5-dsRNA filaments in solution in the presence of 2 mM ADP, which were subsequently adsorbed onto a mica surface. We clearly identified MDA5 monomers on dsRNA (**Fig. S4F**) and can image the complex over time (**Fig. S4G**). In **Video S1**, 6 mM ATP is added at ∼124 s, resulting in translocation and compaction of the MDA5-dsRNA filament (**Fig. 2C**).

To test our model of CARD-dependent filament compaction, we investigate two clinically relevant MDA5 mutants, R338G and I923V, and a mutant without 2CARD (ΔCARD). MDA5 ATPase-dead mutant R338G (**Fig. 1A**) is a pathological gain-of-function mutant that increases signaling and interferon production ^31^. The filaments formed by the R338G MDA5 mutant in the presence of ADP has the same mechanical properties as the ones formed with wild-type (WT) MDA5. They demonstrate similar persistence and contour lengths (**Fig. S5A, Table S7**), underwind coilable dsRNA upon binding (**Fig. S5B**) and has a similar rotation-extension behavior as WT MDA5 (**Fig. S3A** vs. **Fig. S5C**). Removing ADP and excess proteins followed by the addition of 4 mM ATP does neither lead to a decrease of the tether extension (**Fig. 2D**), nor to MDA5 R338G dissociation from the dsRNA tether. Indeed, the analysis of the subsequent force-extension experiment provides similar persistence and contour lengths as preceding ATP addition (inset **Fig. 2D, Fig. S5A, Table S7**). ATP hydrolysis is therefore essential for either MDA5 filament compaction or dissociation.

We next investigate the hyperactive ATPase variant I923V (**Fig. 1A**). MDA5 I923V has a 2-fold greater ATPase activity ^52^ and a loss of antiviral signaling activity relative to WT MDA5 ^11,52,53^. The MDA5 I923V filaments has similar persistence and contour lengths as WT MDA5 filament (**Fig. S5D, Table S7**) and underwind dsRNA upon binding on a coilable tether (**Fig. S5E**). However, rotating the magnetic bead from negative to positive turns results in a complete dissociation of the I923V mutant filament (**Fig. S5F**), indicating that this filament is unstable under torsional stress. Interestingly, addition of either low or high ATP concentration results in two very different responses from the I923V MDA5 filament (**Fig. 2EF**). At low ATP concentration (0.5 mM), we observe the dissociation of MDA5 from dsRNA (**Fig. 2E**), determined via a subsequent force-extension measurement (inset **Fig. 2E, Table S7**). However, at high ATP concentration (4 mM), the filament compacts into a stable and tight complex (**Fig. 2F**), similarly to WT MDA5 filament (**Fig. 2B**). Together, these results support a two-step ATP dependent model for compaction, e.g. MDA5 binding is destabilized by ATP hydrolysis ^8,9^, followed by either rapid binding of another ATP molecule to enable the next hydrolysis cycle or MDA5 dissociation from dsRNA. A mutant such as MDA5 I923V therefore requires higher ATP concentration to compensate the binding instability due to the mutation.

Finally, we investigate how 2CARD impacts the filament compaction with the MDA5 ΔCARD mutant. 2CARD constitutes the N-terminus of MDA5 (**Fig. 1A**) and forms stable interactions with the CARD of MAVS. This association is essential to promote immune response signaling ^54^. Furthermore, 2CARD has been proposed to play another important role in connecting neighboring MDA5’s within the filament ^14^. MDA5 ΔCARD forms a filament on dsRNA with comparable *L*_*C*_ and *L*_*p*_ as the WT MDA5 filament (**Fig. S5G, Table S7**) ^12^. Association of MDA5 ΔCARD to a coilable dsRNA leads to underwinding, similarly to WT MDA5 (**Fig. S5H**). Interestingly, rotation-extension experiments on a filament formed on a non-coilable dsRNA show no change in extension in the absence of ATP, unlike WT MDA5 (**Fig. S5I**). This result suggests that 2CARD bridges consecutive MDA5 protomers in the filament, even in absence of ATP hydrolysis, rendering an initially non-coilable dsRNA tether coilable (see **Fig. S3D** with WT MDA5). Upon addition of ATP into the flow chamber, MDA5 ΔCARD dissociates from dsRNA (**Fig. 2G**), as confirmed by the consecutive force-extension experiment (inset **Fig. 2G, Table S7**). Therefore, in addition to stabilizing MDA5 filaments by connecting adjacent (or nearby) protomers, we have identified a new role for 2CARD in dsRNA compaction.

### MDA5 filaments formation precedes ATP-dependent compaction

All experiments until now have been performed in two steps: filament formation in absence of ATP, followed by flushing out all excess proteins and injection of reaction buffer with ATP. While these experiments have helped to separately monitor filament assembly and ATP-hydrolysis dependent compaction, such conditions are not physiological. We therefore perform experiments where we monitor MDA5 filament assembly and compaction with ATP continuously present (**Fig. 3A-C, Fig. S6A-C**). We investigate only non-coilable dsRNA to circumvent potentially misinterpreting compaction events as the result of dsRNA supercoiling upon MDA5 binding (**Fig. 1GH**).

**Fig. 3:**
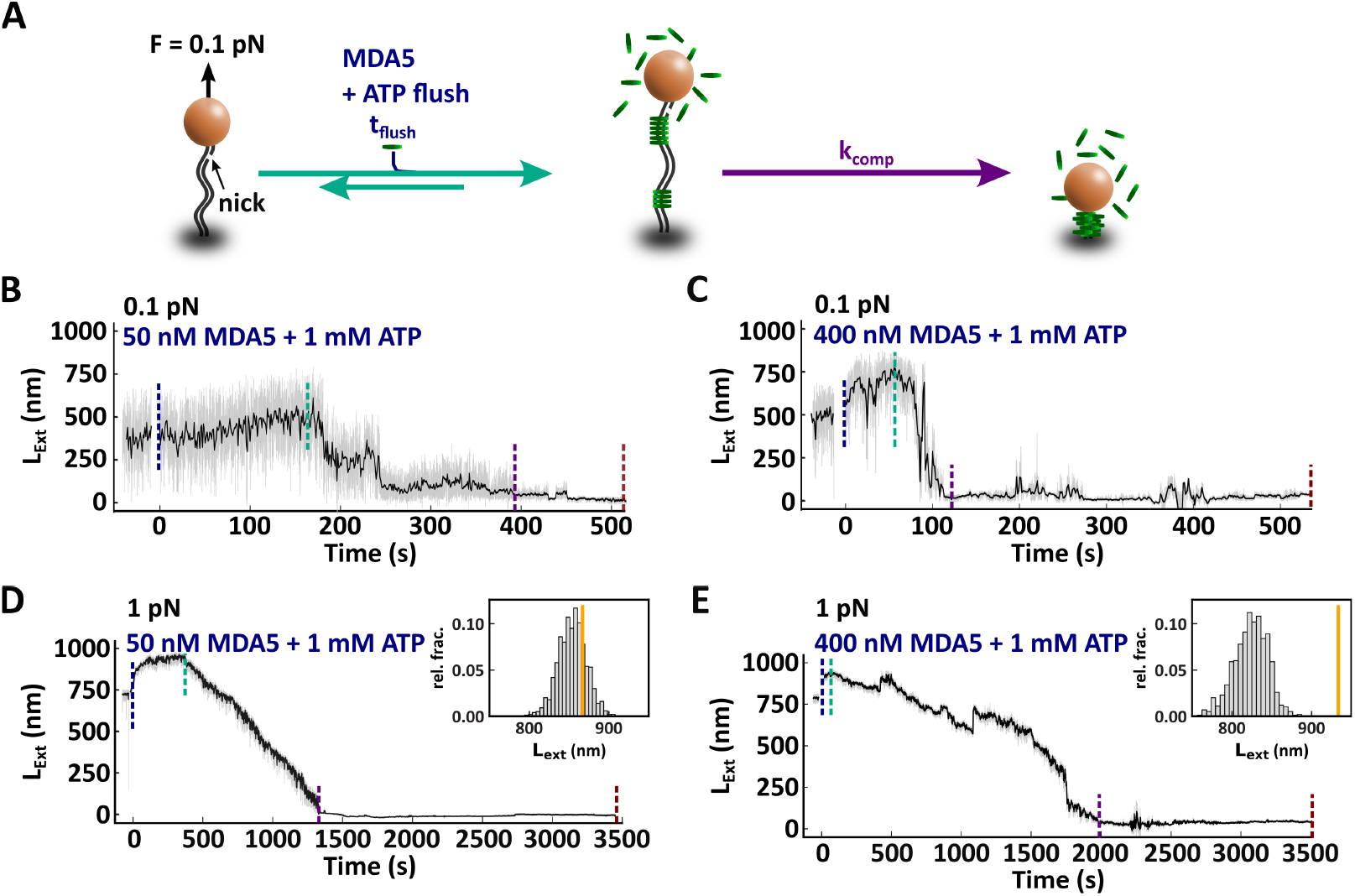
Complete MDA5 filament formation is not required for compaction to occur. **(A)** Description of the experiment monitoring the filament formation upon the direct injection of MDA5 with 1 mM ATP. **(B, C)** Time trace of the filament formation with **(B)** 50 nM or **(C)** 400 nM MDA5 and compaction for the experiment described in (A) at 0.1 pN. **(D, E)** Time trace of the filament formation with **(D)** 50 nM or **(E)** 400 nM MDA5 and compaction at 1 pN. Inset: Relative fraction of the extension of the MDA5-dsRNA filament before compaction. The orange line indicates the mean extension of 50 nM MDA5 and 300 nM MDA5 with 2 mM AMP-PNP at 1 pN as determined by force-extension (**Fig. S1HN**). The vertical dashed lines indicate the end of each phase: flushing (blue), pre-compaction (teal), compaction (purple) and the compacted filament (dark red). The raw (58 Hz) and time averaged (1 Hz) traces are represented in grey and black, respectively.

At low MDA5 concentration (50 nM) and low force (0.1 pN), the filament slowly nucleates (**Fig. 3B**) and starts to form a filament, as noticeable by the increase in the tether’s extension (**Fig. 3B, Fig. S6B**). The extension first increases by ∼100 nm and thereafter decreases by abrupt steps. At such a low force, long distance CARD-CARD interactions of MDA5 monomers bound far away on the dsRNA tether are very likely, explaining the abrupt compaction-decompaction events we observe in the traces (**Fig. 3, Fig. S4D**).

To prevent thermally activated long distance CARD-CARD interactions, we performed the same experiments at higher force (1 pN) (**Fig. 3DE**). At low MDA5 concentration (50 nM), the tether started to extend during the flush of MDA5 until compaction started (∼400 s) (**Fig. 3D**). The extension reached in presence of ATP is similar to the one measured in presence of AMP-PNP (**Fig. S1H, Fig. 3D**, inset). At higher MDA5 concentration (400 nM), we noticed that the filaments extended and thereafter compacted during the flush (**Fig. 3E**), likely due to the cooperative nature of the filament assembly. This results in an extension shorter than a full filament formed with AMP-PNP at 1pN (**Fig. S1N, Fig. 3E**, inset). In conclusion, in the presence of ATP MDA5 filaments forms a first filament before compacting the dsRNA. Even though long-distance CARD-CARD interactions facilitate compaction, they are not required.

### MDA5 filament compaction does not originate from supercoiling

MDA5 filament assembly renders a nicked, and therefore non-coilable, dsRNA coilable (**Fig. S3D**). Structural studies support a change in dsRNA twist induced by MDA5 during the ATP hydrolysis cycle ^8,9^. Therefore, the filament compaction could also originate from plectoneme formation due to a change in dsRNA twist ^36^. To test this hypothesis, we have inserted a 30 nt ssRNA gap near either the 3’-end or 5’-end in one of the two RNA strands or both to rule out any bias in gap orientation (**Fig. 4A, Materials and Methods**). As MDA5 does not bind to ssRNA ^55^, these constructs allow for two separate filament fragments to form (on either side of the gap). Experiments are performed at 1 pN to prevent thermally activated long distance CARD-CARD interactions initiating compaction. In the presence of MDA5 and 1 mM ATP, a fraction of the gapped constructs shows a similar behavior as with the fully dsRNA tether (**Fig. 3D**), i.e. a tether compaction into a stable and tightly compacted form (**Fig. 4B**). This has been verified by the subsequent force-extension experiment (**Fig. 4B**, inset).

**Fig. 4:**
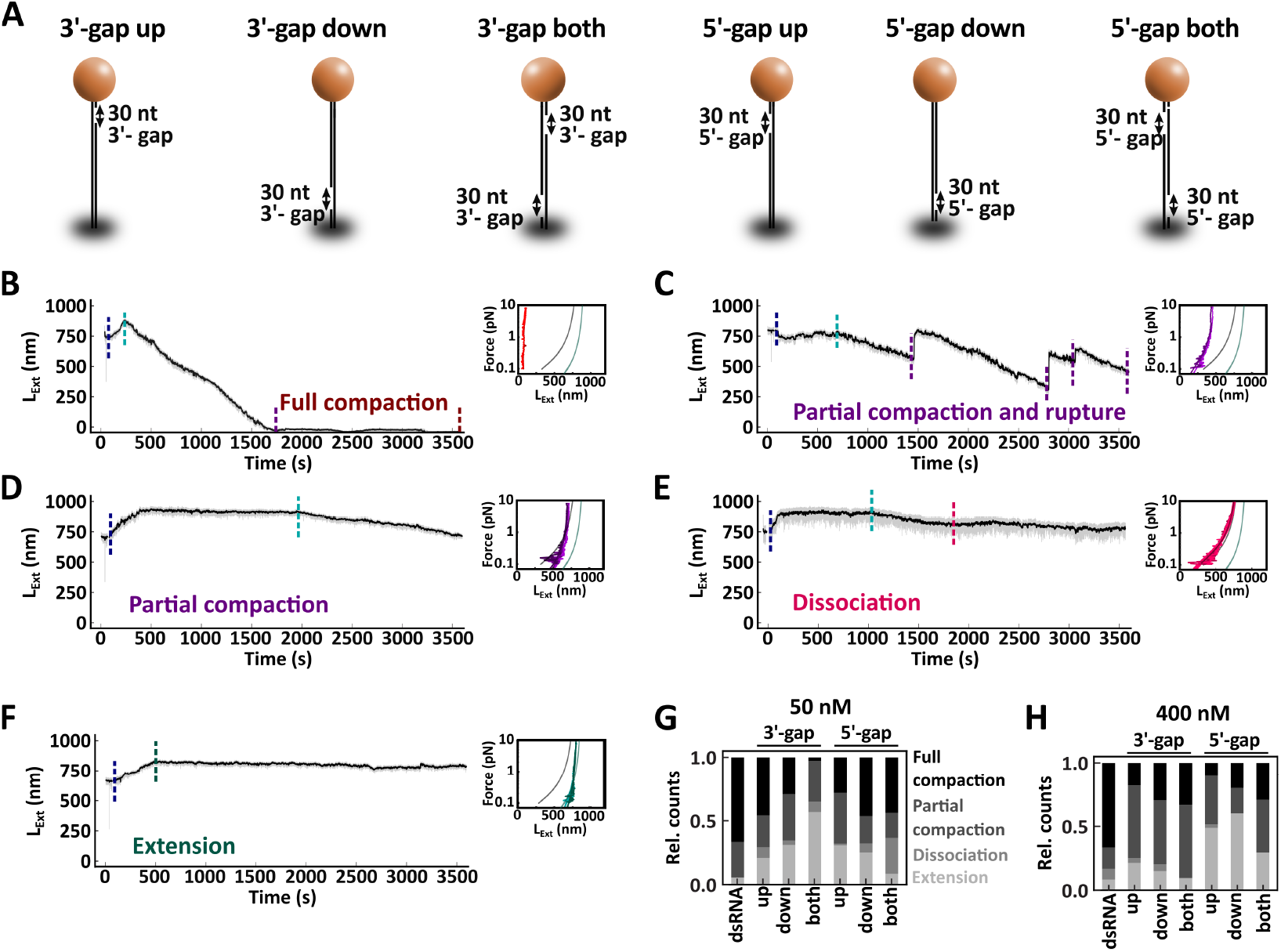
MDA5 compacts constructs with 30 nt gap. **(A)** Description of the dsRNA constructs with 30 nt ssRNA gaps. **(B-F)** Time trace of MDA5 and ATP at 1 pN on gapped dsRNA tether. The tethers can fully compact **(B)**, partially compact with rupture **(C)**, partially compact **(D)**, dissociate **(E)** or fully compact **(F)**. Small inserts: force-extension traces of filament resulting in either extension (green), dissociation (pink), partial compaction (purple), or full compaction (red). Non-extensible WLC fits to force-extension experiments for dsRNA (grey line) and MDA5 filament (teal line). The raw (58 Hz) and time averaged (1 Hz) traces are represented in grey and black, respectively. **(G)** Relative count of compaction events observed on gapped dsRNA with 50 nM MDA5 and 1 mM ATP at 1 pN. **(H)** Relative count of compaction events observed on gapped dsRNA with 400 nM MDA5 and 1 mM ATP at 1 pN. Statistics and mean values provided in **Tables S8 and S9**.

Unlike with fully dsRNA, we also notice partial compaction and rupture (**Fig. 4C**), slow partial compaction (**Fig. 4D**), dissociation (**Fig. 4E**), and tethers that present full and stable extension (**Fig. 4F**). The relative fraction of these events varies with the gap location and orientation, and the concentration of MDA5 (**Fig. 4GH**). At either low or high MDA5 concentration, almost all the fully double-stranded tethers, i.e. without gap, compact. Having a gap in the construct, at either end of either strand led to a lower fraction of full or partial compaction (**Fig. 4GH, Table S8, Table S10**). While increasing MDA5 concentration partially recovers the compacted fraction (full and partial) for the 3’-gap constructs, the 5’-gap ones still result in a smaller fraction (**Fig. 4GH, Table S8, Table S9**). We conclude that MDA5-dependent dsRNA supercoiling cannot explain the tether compaction we observed.

### MDA5 compacts the tether through a dsRNA unwinding and ssRNA loop extrusion mechanism

Finally, we interrogate the mechanism by which MDA5 compacts the dsRNA tether. From the gapped dsRNA constructs experiments (**Fig. 4**), we know that compaction is not due to supercoiling. We hypothesize that compaction occurs after (partial) MDA5 filament assembly through dsRNA unwinding and ssRNA loop extrusion. These loops would therefore be accessible for enzymes that specifically degrade ssRNA, such as RNase A.

To test this hypothesis, we partially assemble MDA5 filaments on coilable dsRNAs at 1 pN and with ADP. We use large magnetic beads (M270, **Materials and Methods**) to apply enough force to break the tightly compacted ribonucleoprotein complex (**Fig. 2B**, inset). The excess proteins are subsequently flushed out and the magnets removed to let the tether relax any torsional stress (**Fig. 5A**). The magnets are then mounted again, and their height adjusted to apply 1 pN force, followed by ATP addition. Once the tethers are fully compacted, the flow cell is treated with RNase A and subsequently rinsed with reaction buffer without proteins. High force is applied to break apart the compacted complex followed by a rotation-extension experiment (**Fig. 5AB**). The addition of RNase A to a previously compacted MDA5 filament renders the tether non-coilable, confirming the partial conversion of the dsRNA tether into ssRNA following ATP-dependent compaction (**Fig. 5B, Table S10**). In absence of either the compaction step, i.e. without ATP (**Fig. 5C, Table S10**), or RNAse A (**Fig. 5D, Table S11**), the tethers remain coilable (**Fig. 5E**). The similar fraction of tethers lost in all experiments likely results from the force jump used to break the compacted filaments (**Fig. 5AE**). AFM investigation further supports that MDA5 can convert dsRNA (**Fig. S7A**) into ssRNA (**Fig. S7B, Video S2**) in the presence of ATP and when no obstacle is present at either 5’-end of the dsRNA.

**Fig. 5:**
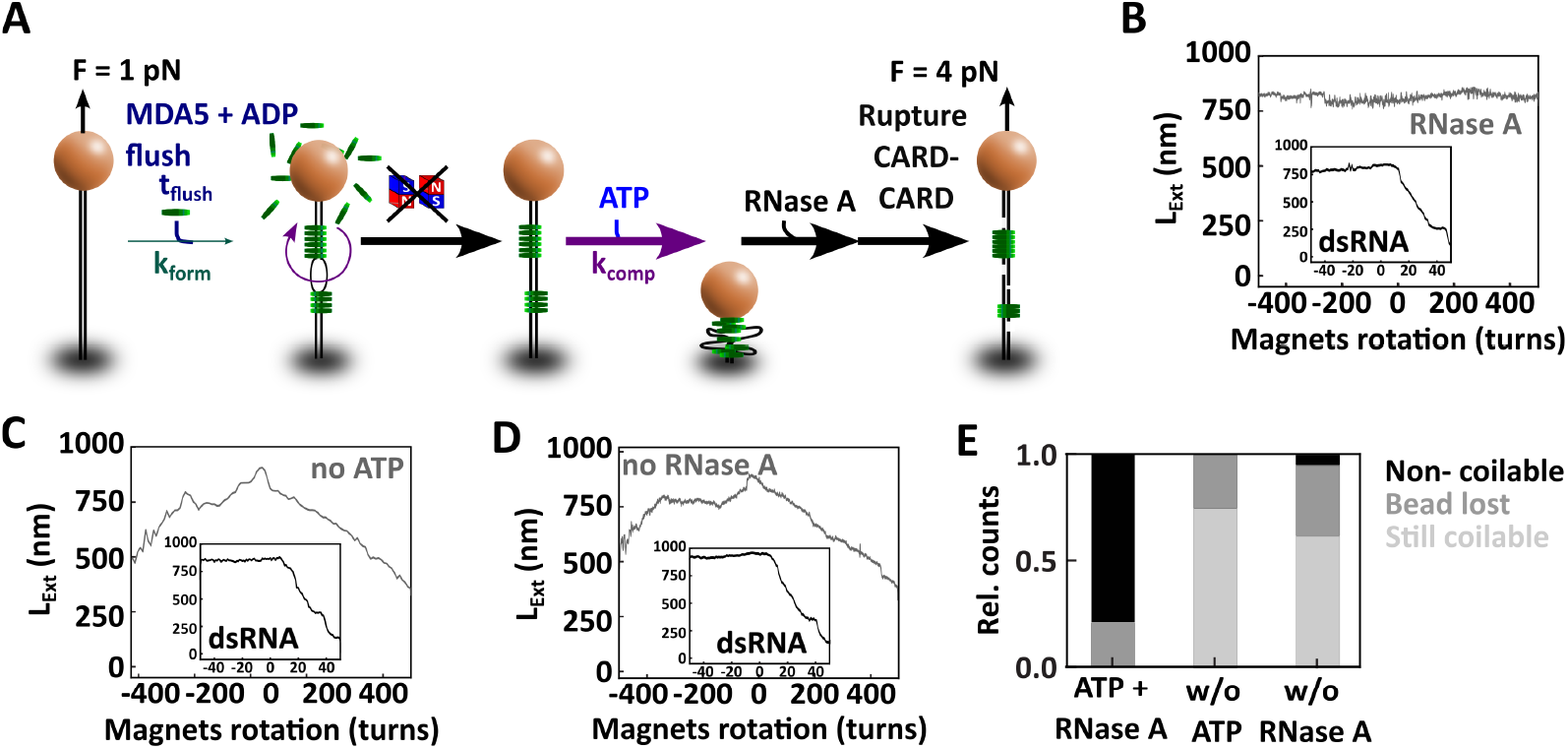
MDA5-RNA complex compaction results from ATP-dependent dsRNA unwinding and single-stranded RNA loops extrusion. **(A)** Schematic of the experiment reporting on ssRNA loop extrusion by MDA5. The filament was formed on a coilable dsRNA with 100 nM MDA5 and 2 mM ADP, the magnets were removed at the end of the assembly to let any torsional stress relax. The free proteins were then flushed out with a reaction buffer containing 4 mM ATP, which led to compaction. Once the MDA5-RNA complex had compacted, RNase A was added to the compacted MDA5-RNA complex. The magnets were thereafter mounted, and high force was applied to break apart the MDA5-RNA complex. A rotation-extension experiment was subsequently performed to evaluate the coilability of the tether. **(B, C, D)** Rotation-extension of an initially coilable dsRNA **(B)** after performing the treatment as described in (A), **(C)** without the ATP addition step described in (A) and **(D)** without the RNase A step described in (A). The insets report on the coilability of the dsRNA at the start of the experiment. **(E)** Relative count of coilable dsRNA that are no longer coilable (black), lost (grey), or still coilable (light grey) at the end of the experiments represented in (B, C, D). Statistics provided in **Tables S10**.

## Discussion

MDA5 plays a crucial role in sensing and recognizing viral dsRNA from the many structured RNAs present in the cell cytoplasm. To achieve this task, MDA5 binds dsRNA and utilizes ATP hydrolysis to proofread dsRNA. Previous biochemical ensemble experiments proposed that ATP hydrolysis promotes dissociation of MDA5 from RNA and that such dissociation is further stimulated by high ATP concentrations ^11,12,14,17^. This raises the question of how MDA5 can recognize dsRNA in the cytosol, where ATP concentration is high, and what is the role of ATP hydrolysis in dsRNA recognition.

To answer this question, we employed single-molecule magnetic tweezers and high-speed AFM to physically manipulate and monitor single dsRNA molecules in interaction with MDA5. We also monitored how wild-type and mutant MDA5s interacted with such a dsRNA in the presence of either ATP or non-hydrolysable analogs. Our data are consistent with MDA5 cooperatively assembling into filaments (**Fig. 1F**) after a slow nucleation step. Nucleation is likely rate-limited by a conformational change, such as accommodating and underwinding dsRNA (**Fig. 1GH**), as it saturates when increasing MDA5 concentration (**Fig. S1C**). In the presence of ATP, MDA5 compacts dsRNA into a tight ribonucleoprotein complex that requires forces as high as ∼20 pN to break apart (**Fig. 2**). Both ATP hydrolysis and CARD-CARD oligomerization are required for the compaction to occur (**Fig. 3**). MDA5 ATPase activity is sufficient to compact dsRNA against opposing forces of up to 4 pN (**Fig. S4A-D**). This further supports that tether compaction does not require thermally activated long-range interactions to occur, even though such interactions accelerate the compaction process in absence of force (**Fig. 3BC**). This may also explain why MDA5 is capable of dislodging proteins tightly bound to dsRNA ^18,19,52^. We showed that compaction does not result from tether supercoiling (**Fig. 4**), but rather through dsRNA unwinding and ssRNA loop extrusion (**Fig. 5**). The presence of ssRNA gaps in the dsRNA construct decreases the fraction of fully compacted dsRNA, indicating that MDA5 senses such gaps and thereafter dissociates (**Fig. 4**).

Based on our experimental results, we propose the following model (**Fig. 6**). After a slow nucleation step, the MDA5 filament assembles cooperatively and directionally, stabilized by CARD-CARD oligomerization that prevents MDA5 dissociation post ATP hydrolysis. Cycles of ATP hydrolysis result in dsRNA compaction. This process goes on until all bare dsRNA has been pulled through the MDA5 filament and reaches the next obstacle. In our assay, such obstacles can be either the magnetic bead and the glass surface attachment points, or another MDA5 filament originating from another nucleation site. In the cellular context, we propose that LGP2 acts as an obstacle for MDA5 translocation when bound to dsRNA internally^19,56^ or at either dsRNA ends ^10,57^ (**Fig. 6**). Indeed, LGP2 bound to dsRNA has been shown to potentiate MDA5 signaling ^19,26,58–61^. In the absence of an obstacle at the dsRNA end, MDA5 unwinds the dsRNA completely and eventually falls off from the free ends (**Fig. S7B, Video S2**). As a result, no compacted ribonucleoprotein complex is produced. Our model is further supported by a recent single-molecule fluorescence study showing that LGP2 blocks MDA5 translocation along dsRNA ^20^. Moreover, in absence of a stretching force, conformational dsRNA chain fluctuations enable long distance CARD-CARD interactions to lock together distant MDA5 monomers on the same or a different dsRNA, which further promotes rapid compaction (**Fig. 2B, Fig. S4A**). We propose that MDA5 helicase activity is impaired once the filament has fully compacted, due to CARD-CARD oligomerization (**Fig. 6**). It remains to be determined whether this tightly compacted form plays a role in signaling. Aggregation of MAVS via CARD-CARD interactions is essential in MDA5 signaling ^13^ and only oligomerized 2CARD can activate MAVS ^14^. Our data shows that CARD-CARD association occurs between neighboring WT MDA5 in absence of ATP hydrolysis, as non-coilable tethers are rendered coilable following full filament assembly with WT MDA5 (**Fig. S3D**), but never with MDA5 ΔCARD (**Fig. S5I**). The stability of the compacted ribonucleoprotein complex (**Fig. 2B**) suggests that this form likely plays an essential role in capturing viral dsRNA, i.e. making them thereby unavailable for viral replication, while potentially ensuring an effective signaling when associated with the MAVS-CARD.

**Fig. 6:**
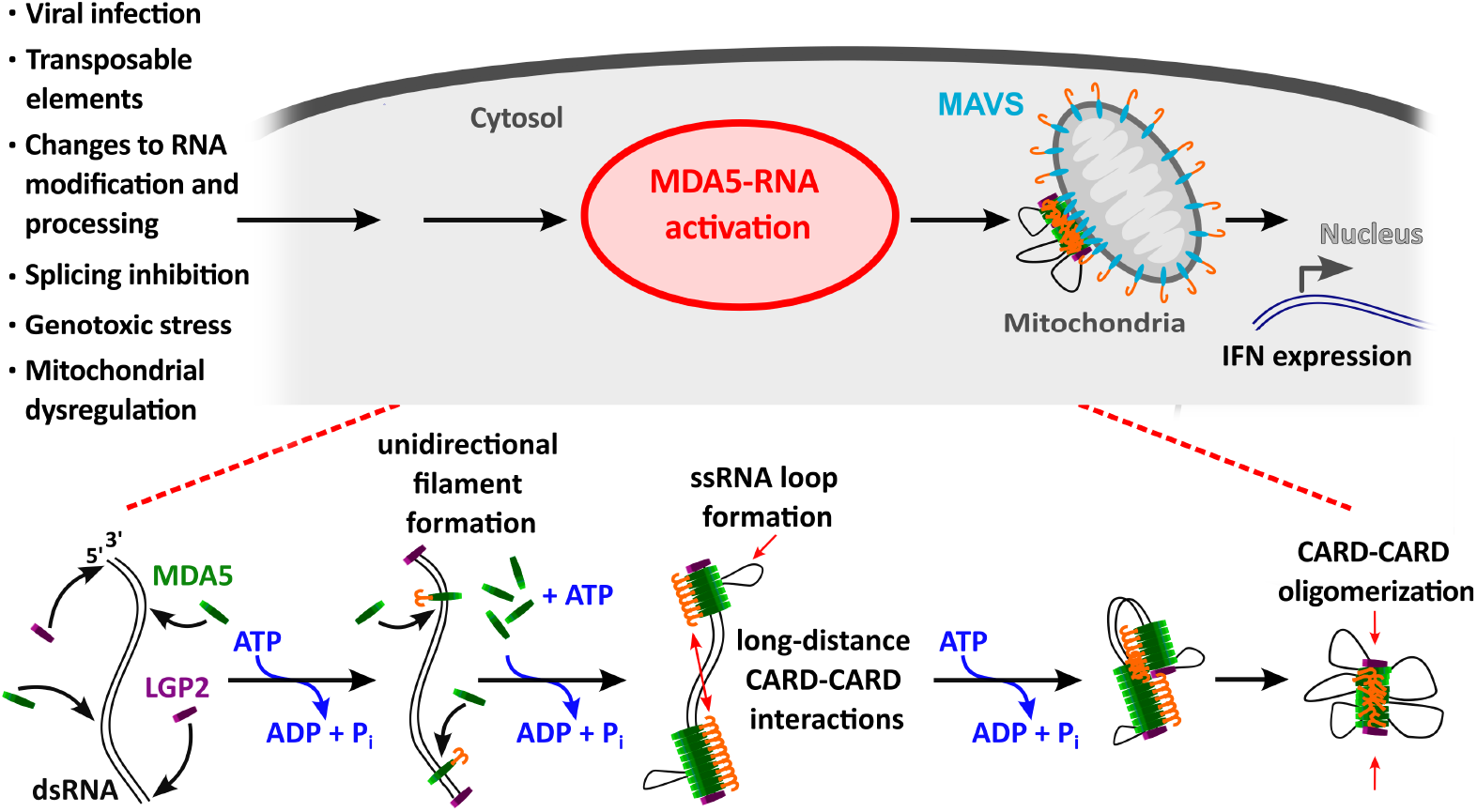
Mechanism of dsRNA compaction by MDA5. The presence of long dsRNA in the cytosol is related to many cellular dysfunctions and viral infections and induces MDA5 activation. MDA5 rapidly loads onto the dsRNA forming filaments cooperatively and unidirectionally. Employing ATP hydrolysis MDA5 unwinds the dsRNA and extrudes ssRNA loops, pulling inwards the dsRNA ends, forming a tightly compacted nucleoprotein complex. Furthermore, ATP hydrolysis exposes the CARD domain, enabling long-distance CARD-CARD interactions and oligomerization, stabilizing the filament on dsRNA. LGP2 associated with the dsRNA prevent the filament to fall off the RNA, thereby stabilizing the nucleoprotein filament into a tight complex. Finally, the compacted MDA5-RNA complex can now associate with the MAVS CARD to enable further signaling towards interferon (IFN) expression.

Our study highlights a novel function for MDA5 ATPase activity, namely to compact dsRNA through ssRNA loop extrusion into a tightly compacted ribonucleoprotein complex stabilized by CARD-CARD oligomerization. We propose that the resulting complex captures viral dsRNA to prevent further usage for viral replication. Future work will explore whether other factors potentiate or antagonize this new function and whether compaction is involved in RNA proofreading.

## Materials and Methods

### dsRNA synthesis

A comprehensive protocol for synthesizing dsRNA constructs for magnetic tweezers has been provided in reference ^38^. Briefly, a 3.2 kbp double-stranded RNA (dsRNA) was assembled from single-stranded RNA (ssRNA) oligomers. DNA fragments from pBAD plasmid were amplified using Phusion polymerase with primers (**Table S11**) that contained the T7 promoter. Handles containing Digoxigenin-UTP (Jena Biosciences) or Biotin-UTP (Jena Biosciences) were also amplified to enable attachment to the surface of the flow cell and the magnetic bead, respectively. In vitro transcription was carried out using the HiScribe kit (NEB) to generate ssRNA fragments. If coilability of the tethers was required, the ssRNA fragments were mono-phosphorylated using pyrophosphatase (Thermo Fisher) before annealing and ligated using T4 RNA ligase 2 (NEB). Constructs containing 30 nt gaps, were constructed via shorter ssRNA oligomers for the required strand (‘up’ or ‘down’ strand) and annealed to the full-length strand with opposite orientation (**Table S12**).

### MDA5 expression and purification

A detailed protocol of the cloning, expression and purification procedure can be found in ^8,9^. MDA5 was cloned into pET28a plasmid with an N-terminal 6x His-tag and TEV protease cleavage site for purification. The MDA5 gene encodes for the mouse *MDA5 (IH1H1)* without residues 646-663 (in this work termed MDA5 WT). Mutants were cloned by overlap PCR. MDA5 WT was expressed in *E*.*coli* BL21(DE3). The lysate was loaded onto Ni-NTA agarose and eluted using a buffer gradient containing imidazole. After a subsequent anion exchange column, the protein was further purified in a size exclusion chromatography.

### Flow cell preparation

A comprehensive protocol for efficient passivation of flow cells to be used in magnetic tweezers experiments is described in detail in reference ^37^. The top coverslip (#1 24 × 60 mm, Menzel GmbH Germany) were drilled with two holes using a sandblaster (Problast 2, Vaniman USA) and Al_2_O_3_ particles (34-82 µm, F230, Eisenwerk Wuerth Germany). Both top and bottom coverslips were sonicated in Hellmanex III (Sigma Aldrich) solution (2% V/V in demineralized water) and the bottom coverslip were functionalized by spreading across its surface 4 µl of 1 mg/ml nitrocellulose dissolved in amyl acetate. The flow cell was assembled by sandwiching between a top and bottom coverslip a double layer of Parafilm M with a carved-out channel to connect the two holes of the top coverslip. The flow cell was sealed by melting the Parafilm on a hot plate at 100°C for ∼2 min.

As reference beads, either 1.1 µm or 3 µm polystyrene beads (Sigma Aldrich) were used when using either MyOne or M270 Dynabeads (Thermo Fisher) magnetic beads, respectively. To functionalize the flow cell, 50 µl of 50 µg/ml anti-digoxigenin (Roche) was applied and incubated for 30 min. Afterwards, the flow cell was rinsed and flushed with 50 µl of 1 mg/ml BSA (NEB) to prevent non-specific binding of the sample.

The dsRNA tethers were mixed with either 1.1 µm or 3 µm superparamagnetic streptavidin coated beads (MyOne or M270 Dynabeads, Thermo Fisher) and incubated for 5 min before being flushed into the flow cell ^38,39^. The unspecific and weakly bound tethers were washed out, and the tethers were further selected for their length and coilability (if applicable).

### Magnetic tweezers experiments

The experiments were conducted using a high-throughput magnetic tweezers instrument, which has been previously described in details in references ^34,37,62–64^. Briefly, the setup consists of an inverted microscope equipped with a 60x oil immersion objective lens (Nikon Plan APO 60X /1.4 NA Oil) and a CMOS camera (Dalsa Falcon 2 FA-80-12 (12 Mpixel)). The position of the objective can be adjusted using a PIFOC piezo (P-726, Physik Instrumente Germany). An LED (λ=660 nm, 400 mW, LH CP7P, Lumitronix) is used to illuminate the flow cell, which is mounted on top of the objective and equipped with an aspheric condenser lens (ACL25416U-A, ø1”, f=16 mm, Thorlabs). Two vertically aligned 5 mm permanent magnets (SuperMagnete, Germany) are mounted on top of the flow cell and can be translated in the z-direction and rotated using linear motors (M-126-PD1 and C-150, respectively, Physik Instrumente Germany) as required. The beads tridimensional position are tracked in real-time using the bead tracker described in reference ^40^. The software and its implementation in LabView are publicly available at https://github.com/jcnossen/qtrk and https://github.com/jcnossen/BeadTracker, respectively. All experiments were performed using a 17 ms shutter time and a 58 Hz acquisition frequency. All time traces presented in this work were filtered by a moving average filter with 58 frames window size.

The MDA5 protein aliquots were stored at a concentration of 1-15.7 µM at −80°C until use. Prior to experiments, the aliquots were thawed and equilibrated with 2 mM ADP at 37°C for 90 minutes if applicable. All experiments were conducted in MDA5 measurement buffer (20 mM HEPES, 40 mM sodium chloride, 40 mM potassium chloride, 5 mM magnesium chloride, 2 mM sodium azide, 1 mg/mL BSA, and 1 mM DTT) unless otherwise specified.

We used 3.2 kbp dsRNA to tether the magnetic beads to the flow cell surface. To initiate filament formation, MDA5 and nucleotide were mixed prior to be flushed into the flow cell. Filament formation experiments were performed in the presence of 2 mM ADP at a force of 0.1 pN, unless stated otherwise.

Compaction experiments with pre-assembled MDA5 filaments in the presence of either ADP or AMP-PNP were performed by flushing 200 µl reaction buffer containing ATP into the flow cell after the filament assembly, flushing out excess proteins.

Rotation extension experiments were conducted to determine the coilability of the dsRNA tether at forces of 0.5 pN and 4 pN, with rotations ranging from −50 to +50 turns before the addition of MDA5. After filament formation, the rotation extension was carried out at the indicated force, with rotations ranging from −500 to +500 turns.

### High-speed Atomic Force Microscopy

The High-Speed Atomic Force Microscopy (HS-AFM) videos were generated using amplitude modulation tapping mode imaging in liquid (RIBM Japan), using USC-F1.2-k0.15 cantilevers (NanoWorld, Switzerland) with a nominal spring constant of 0.15 N/m and a resonance frequency ≈ 0.6 MHz in liquid ^51,65^. All HS-AFM recordings were performed at room temperature and in reaction buffer (20 mM HEPES pH 7.9, 40 mM sodium chloride, 40 mM potassium chloride, 5 mM magnesium chloride, 2 mM sodium azide). The dsRNA (with or without proteins) was incubated using the measurement buffer prior to imaging. A minimal imaging force was maintained by using ∼1 nm cantilever-free amplitude and ∼0.8 nm set point amplitude. To follow the MDA5-dsRNA interactions, a mixture of dsRNA, 50 nM MDA5, and 2 mM ADP were prepared 15 minutes prior to surface binding. After incubation of the above-mentioned mixture for 15 minutes on mica, HS-AFM imaging was conducted on a liquid chamber containing 40 µl of measurement buffer. Once a dsRNA-MDA5 complex was localized by HS-AFM imaging, a 10 µl ATP stock was added to the chamber to reach a final ATP concentration of 6 mM. Images were primarily processed using built-in scripts (RIBM, Japan) in Igor Pro (Wavemetrics, Lake Oswego, OR, USA) and analyzed using ImageJ software.

### Fitting of force extension data

Force-extension experiments for forces below 8 pN were performed using MyOne magnetic beads and repeated three successive times on each tether before and after the filament has formed and equilibrated. The magnets were lowered at a rate of 0.1 mm/s to progressively increase the force on the tether from 0.1 to 8 pN.

To extract the mechanical properties of the tether, i.e. persistence length *L*_*p*_ and contour length *L*_*C*_, a non-extensible worm-like chain model ^46^ has been fitted to the individual force-extension trace:

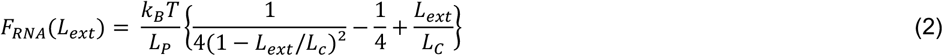

with Boltzmann constant *k*_*B*_, temperature *T* and tether extension *L*_*Ext*_.

However, large fluctuations caused by Brownian motion can affect the accuracy of position estimation ^41^. To overcome this issue, we integrated the force-extension trace to collapse the Brownian motion onto the function:

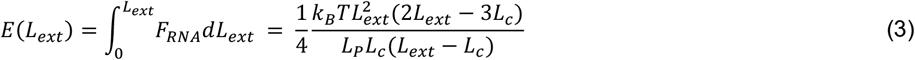

We only considered 0 *nm* ≤ *L*_*p*_ ≤ 500 *nm* and 7--00 *nm* ≤ *L*_*C*_ ≤ 2000 *nm* as acceptable fitting results. The mean and standard deviation of *L*_*p*_ and *L*_*C*_ were extracted from fitting the individual traces using **Equation 3** and used to derive the mean force-extension trace. The worm-like chain model (**Equation 2**) represented in this manuscript use *L*_*p*_ and *L*_*C*_ calculated from **Equation 3**.

Force-extension experiments for forces up to 50 pN were performed using M270 magnetic beads. The individual tethers were stretched from 0.5 pN to 50 pN by moving the magnets towards the flow cell at 0.01 mm/s. The force was subsequently lowered at the same magnets translation rate. Only one cycle on the tether before and after addition of the protein was performed. Rupture forces of compacted MDA5-dsRNA filaments were selected manually from individual traces.

### Extracting the filament nucleation, formation, translocation and compaction rates

To determine the filament nucleation rate *k*_*nucl*_ we manually selected on the 1 Hz low-pass filtered traces the time between the start of the flush of MDA5 in the flow cell and the time where the tether extension increased (**Fig. 1C**).

Concerning filament formation and compaction rates *k*_*form*_ and *k*_*comp*_, we first manually selected the regions in the trace where the tether extension either increased or decreased, respectively. *k*_*form*_ and *k*_*comp*_ were subsequently extracted using a sliding average window of 20 seconds and 50 seconds, respectively, in the selected regions. Only compaction rates larger than 0.1 nm/s (detection limit) were considered. For *k*_*form*_, only increases larger than two times the variance in the trace signal before the MDA5 flush were considered. Filament formation rates were normalized to the footprint of an MDA5 monomer according to

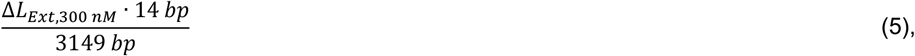

where Δ*L*_*Ext*,300 *nM*_ is the change in extension at 0.1 pN upon filament formation with 300 nM WT MDA5 and 2 mM AMP-PNP, 14 bp being the footprint of WT MDA5 and 3149 bp is the length of the dsRNA tethers used in this assay.

MDA5 translocation rates were determined by dividing the length of the dsRNA (3149 bp) by the time between the addition of ATP to a formed MDA5 filament and the end of dissociation of MDA5 from the filament, i.e. the point at which the extension of the tether represents bare dsRNA (indicated as “dissociation” in **Fig. 5G**).

For the detection of rupture events during filament compaction (**Fig. S4E**), a threshold of twice the standard deviation of the raw signal before addition of ATP was defined. If the extension of the averaged signal crossed this threshold within a 5 second time window, a rupture event was counted.

Time traces of MDA5 in presence of 1 mM ATP at 1 pN were classified according to their extension during the experiment (**Fig. 4**). Tethers that compacted to a stable oligomer that had no force-extension response were classified as ‘fully compacted’. Tethers whose extension decreased below the initial extension of bare dsRNA but did not form a stable oligomer (e.g. because of a rupture event) were classified as ‘partially compacted’. For some traces a decrease in extension was observable but their extension did not drop below the extension of the bare RNA tether. These traces were classified as ‘dissociated’ even if the force-extension measurement indicated a higher persistence length than dsRNA. As it was not possible to distinguish dissociation from very slow compaction, these traces were categorized together. Tethers whose extension did only increase after MDA5 and ATP addition but remained otherwise stable, were classified as ‘extended’ (**Table S8, Table S9**).

### Fitting the Hill equation on the normalized persistence length *L*_***p***_

The normalization of the persistence length *L*_*p*_ was performed as described in reference ^48^:

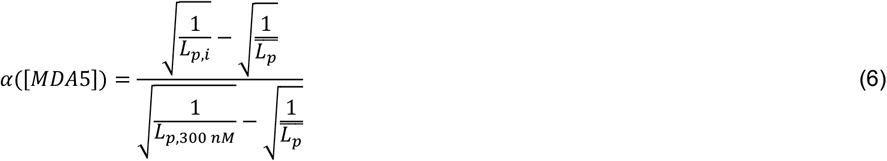

where α([*MDA*5]) is the fractional occupancy of the MDA5-dsRNA filament, *L*_*p,i*_ is the persistence length of the filament at a given MDA5 concentration i, 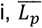 is the mean persistence length, and *L*_*p*,300 *nM*_ is the persistence length at 300 nM MDA5. We assume that a complete filament was formed at 300 nM MDA5.

### Error estimation for fraction of filament events

We calculated the relative fraction *f*_*rel*_ of an event *N*_*i*_ over all events *N*_*all*_ and estimated the error *δf*_*rel*_ according to

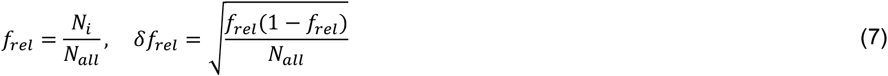

The Hill equation (**Equation 1**) was fit to the fraction occupancy *α*([*MDA*5]) as calculated with **Equation 6** using a non-linear least squares fit with *δf*_*rpl*_ as uncertainty. The fitting was averaged over 1000 bootstrap samples.

## Acknowledgments

We thank the lab of Annemarthe van der Veen for fruitful discussions and Craig E. Cameron for valuable input to the manuscript.

## Funding

DD was supported by the Interdisciplinary Center for Clinical Research (IZKF) at the University Hospital of the University of Erlangen-Nuremberg, “BaSyC – Building a Synthetic Cell” Gravitation grant (024.003.019) of the Netherlands Ministry of Education, Culture and Science (OCW) and the Netherlands Organization for Scientific Research (NWO), and the NWO-M Open Competition Domain Science grant (OCENW.M.21.184). YM and RS were supported by the Wellcome Trust (awards 217191/Z/19/Z and 215378/Z/19/Z, respectively). AHdV was supported by the Human Frontier Science Program (fellowship LT000454/2021-L). WHR was supported through the EU INFRAIA consortium MOSBRI.

## Author contributions

SQ, YM and DD designed the research. SQ, SM, WHR and DD designed the single-molecule experiments. AHdV, RS and YM expressed and purified recombinant MDA5 proteins. QS and FSP provided the RNA construct. SQ and SM performed the single-molecule experiments. SQ, SM and PPBA analyzed the data. CPB provided theoretical support. SQ, SM, MK, WHR and DD interpreted the data. SQ and DD wrote the article. All the authors read and edited the article.

## Competing interests

YM is a consultant for Related Sciences LLC and has profits interests in Danger Bio LLC. None of the other authors have any conflicts of interest with the content of this article.

## Data and materials availability

The data of this study are available from the lead author.

## Notes

### Competing Interest Statement

Yorgo Modis is a consultant for Related Sciences LLC and has profits interests in Danger Bio LLC.

### Summary of Updates

Additional experiments were performed to further support the conclusions of the paper. These experiments, their analysis and descriptions were added to the manuscript.

