## Supplementary Information for "MDA5 generates compact ribonucleoprotein complexes via ATP-dependent single-stranded RNA loop extrusion"

Salina Quack *et al.*

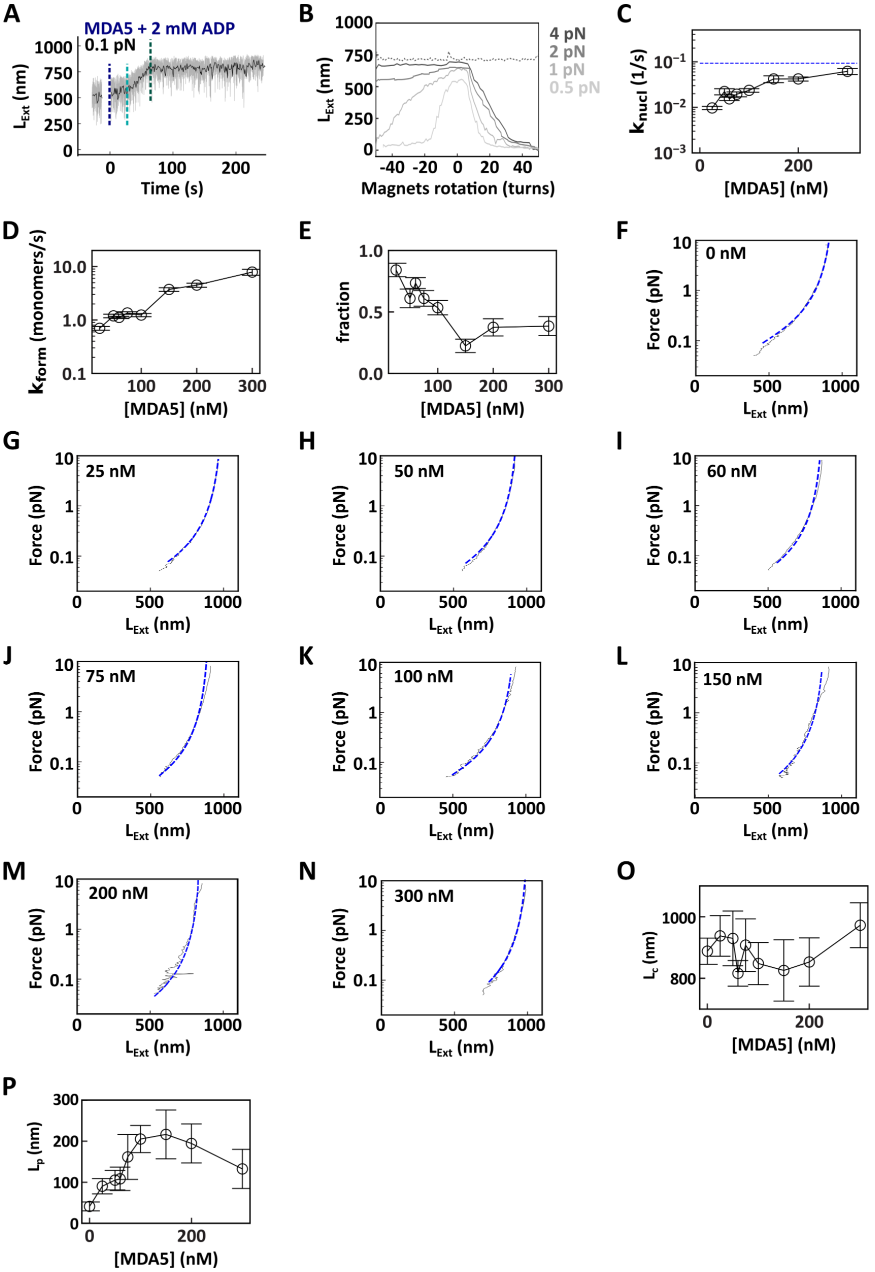


**Figure S1: Force-extension traces of MDA5 filaments at different MDA5 concentrations. (A)** Filament formation trace of 100 nM MDA5 and 2 mM ADP at 0.1 pN experimental conditions. The raw (58 Hz) and time averaged (1 Hz) traces are represented in grey and black, respectively. The vertical dashed lines indicate the end of each phase: flushing of MDA5 with ADP (dark blue), nucleation (teal), and filament formation (green). **(B)** Rotation-extension traces of non-coilable dsRNA (dotted line) and coilable dsRNA (solid lines) at various forces. **(C)** Mean nucleation rate of MDA5 on dsRNA in the presence of AMP-PNP. The blue dotted line indicates the limit for the nucleation rate based on the time it takes to add the proteins to the flow cell. **(D)** Mean filament formation rate on dsRNA in the presence of AMP-PNP. **(E)** Relative fraction of nucleation events that were longer than the flush duration. **(F- N)** Mean force extension (black line) and a fit using the worm-like chain (WLC) model (blue dashed line, **Equation 2**). Force extension of **(F)** ~3.2 kbp dsRNA and MDA5 filaments formed using either **(G)** 25 nM, **(H)** 50 nM, **(I)** 60 nM, **(J)** 75 nM, **(K)** 100 nM, **(L)** 150 nM, **(M)** 200 nM, or **(N)** 300 nM WT MDA5, respectively, in the presence of AMP-PNP. Statistics and WLC model fits parameters values are provided in **Table S2**. **(O)** Mean contour length and **(P)** persistence length of MDA5 filaments. The error bars are one standard deviation. (C, D, O, P) Error bars represent the standard deviation from 1000 bootstraps. The lines are guides for the eye. Statistics, mean, and error values are provided in **Table S1** and **Table S2**.

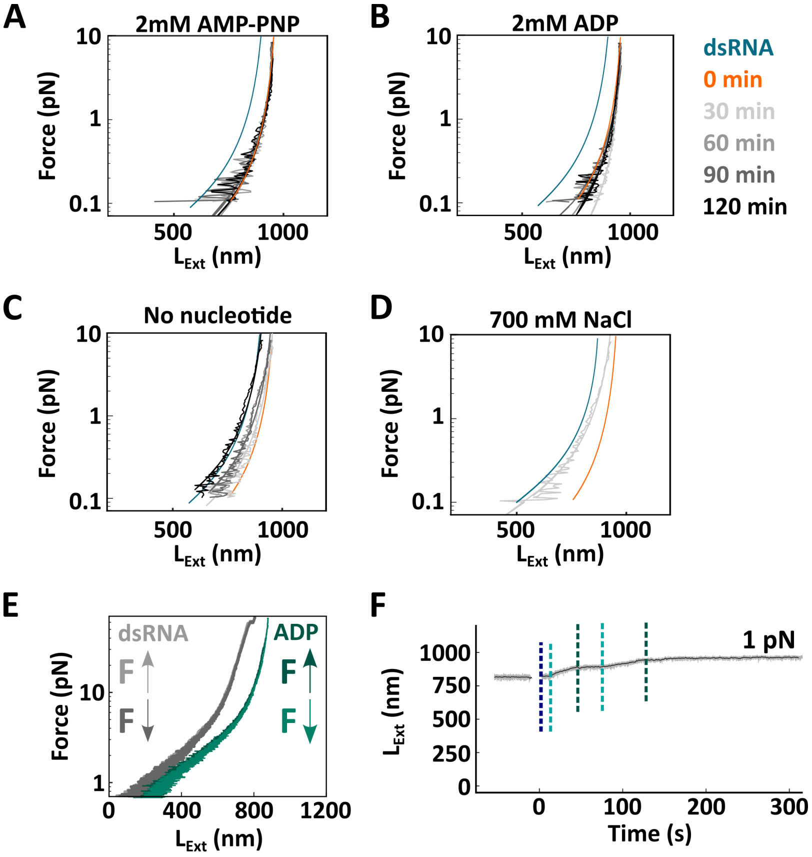


**Figure S2: MDA5 filament stability in different conditions, pauses in filament assembly and assembly kinetics on a coilable dsRNA tether.** Mean force extension of MDA5 filament in the presence of either **(A)** 2 mM AMP-PNP or **(B)** 2 mM ADP, **(C)** formed in absence nucleotide, or **(D)** after treatment with the standard reaction buffer containing 700 mM NaCl in comparison to bare dsRNA (blue) and a MDA5 filament at t=0 min (orange). Color code described on the right-hand side of panel (B). Statistics, mean, and error values from the WLC fits are provided in **Table S3**. **(E)** Force-extension of dsRNA (grey) and MDA5 filament (green) in the presence of ADP upon dynamically increasing (light) and decreasing force (dark). **(F)** Filament formation trace of MDA5 in the presence of AMP-PNP on a non-coilable dsRNA tether at 1 pN.

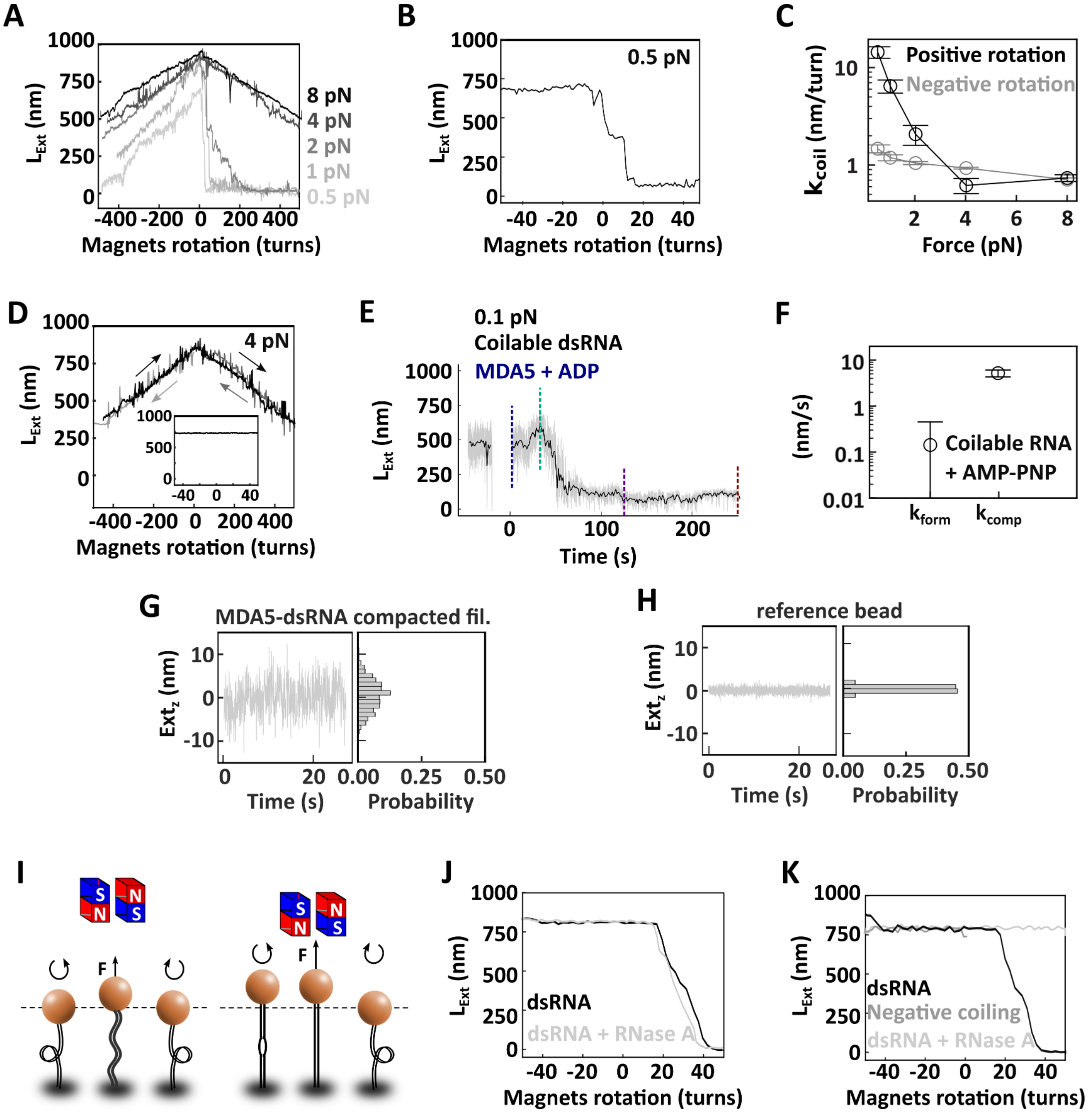


**Figure S3: Torsional behavior of the MDA5-dsRNA filament. (A)** Rotation extension of MDA5-dsRNA filaments in presence of ADP at 0.5 pN (light grey), 1 pN (grey), 2 pN (medium grey), 4 pN (dark grey), and 8 pN (black). **(B)** Rotation-extension trace of MDA5 filament in the presence of 2 mM ADP between -50 and 50 turns. **(C)** Mean decrease in extension rate from data in (F) for either negative (grey) or positive (black) turn addition. The error bars are the standard deviation from 1000 bootstrap samples. The lines are guides for the eye. Statistics, mean, and error values are provided in **Table S4**. **(D)** Rotation-extension of a non-coilable dsRNA tether after nucleoprotein filament formation with MDA5 and ADP. The arrows indicate the direction of rotations applied. Insert: Rotation extension trace of the tether at 4 pN before MDA5 injection in the flow chamber. **(E)** Time trace of a filament assembly on a coilable dsRNA. The vertical dashed lines indicate the end of each phase: flushing of 100 nM MDA5 with 2 mM ADP (blue), respectively, nucleation (teal), filament formation (green), filament compaction (purple) and the compacted filament (red). The raw (58 Hz) and time averaged (1 Hz) traces are represented in grey and black, respectively. **(F)** Mean formation and compaction rates of MDA5 filament on a coilable dsRNA tether. Error bars represent the standard deviation from 1000 bootstrap samples. Statistics, mean, and error values are provided in **Table S5. (G)** Time trace of an MDA5-dsRNA filament after compaction with probability histogram of the fluctuations in the z-extension. **(H)** Time trace of a reference bead. with probability histogram of the fluctuations in the z-extension. (G, H) The raw (58 Hz) and time averaged (1 Hz) traces are represented in grey and black, respectively. **(I)** Description of a rotation-extension experiment of a coilable dsRNA tether at low and high force (left and right, respectively). **(J)** Rotation-extension trace of a coilable dsRNA tether at 4 pN before (black) and after (light grey) the addition of 10 ng RNase A while the tether was torsionally relaxed and at 4 pN. **(K)** Rotation-extension trace of the same dsRNA tether at 4 pN before (black) and after (light grey) the addition of 10 ng RNase A while the tether was negatively supercoiled and at 4 pN.

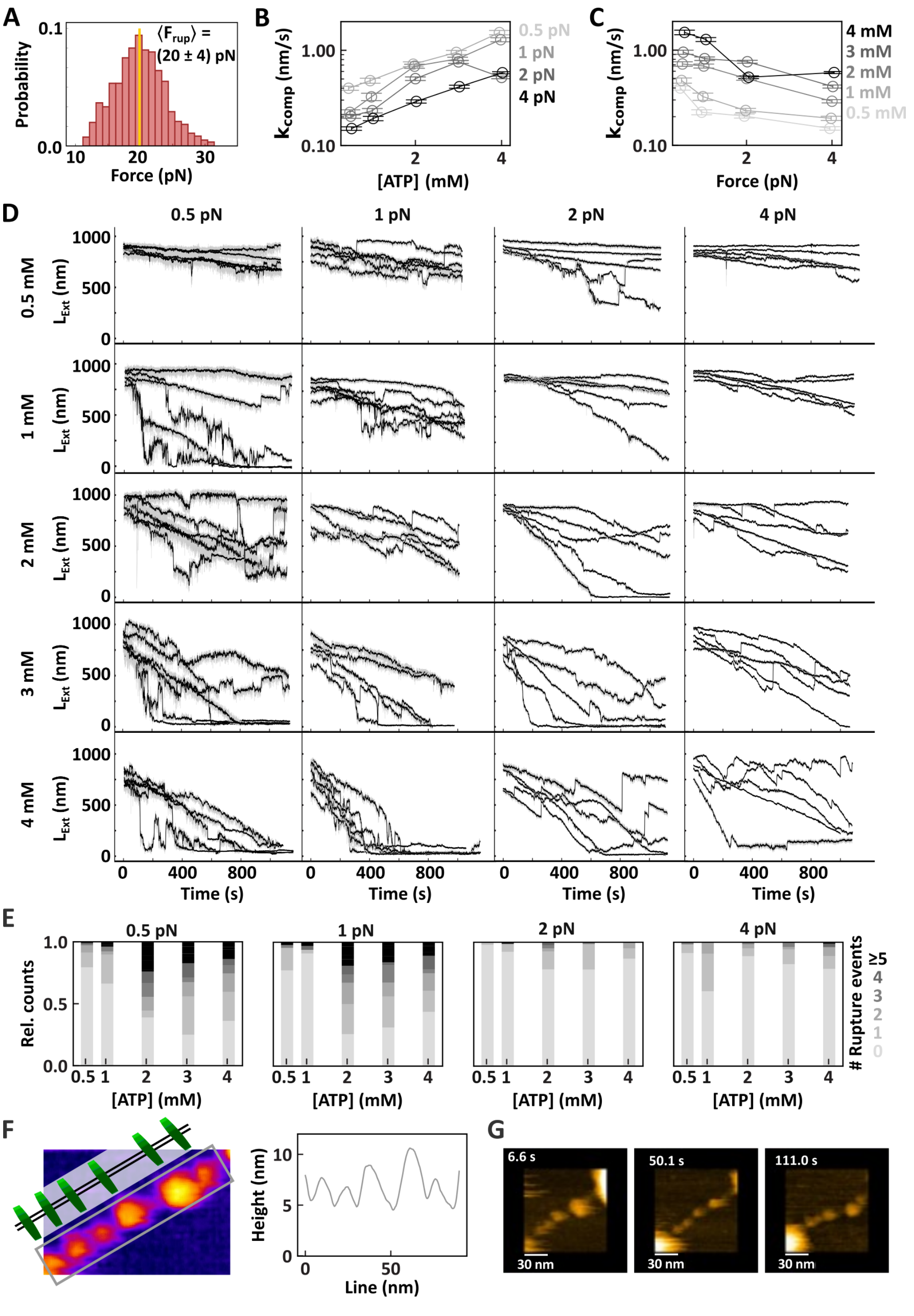


**Figure S4: MDA5 filament compaction is force and ATP concentration dependent. (A)** Distribution of rupture forces observed when breaking the oligomerized filament after ATP hydrolysis. Mean rupture force indicated in yellow and above the plot (the error is one standard deviation). **(B, C)** Mean compaction rate of MDA5-dsRNA nucleoprotein filaments for the **(B)** force or **(C)** ATP concentration indicated above. **(D)** Time traces of filament compaction in the presence of the indicated concentration of ATP and force. The raw (58 Hz) and time averaged (1 Hz) traces are represented in grey and black, respectively. **(E)** Relative counts of rupture events for filament compaction experiments as shown in (A). **(F)** AFM image of dsRNA-MDA5 complex in presence of 2mM ADP (left) and schematic of the MDA5-dsRNA filament. Height profile (right) along a line cross section through the middle of the grey rectangle (left). **(G)** Snapshot HS-AFM images of the dsRNA-MDA5 complex in the presence of 2 mM ADP imaged over time. The imaging rate is 300 milliseconds per frame. **(B, C, E)** Statistics, mean, and error values are provided in **Table S6**.

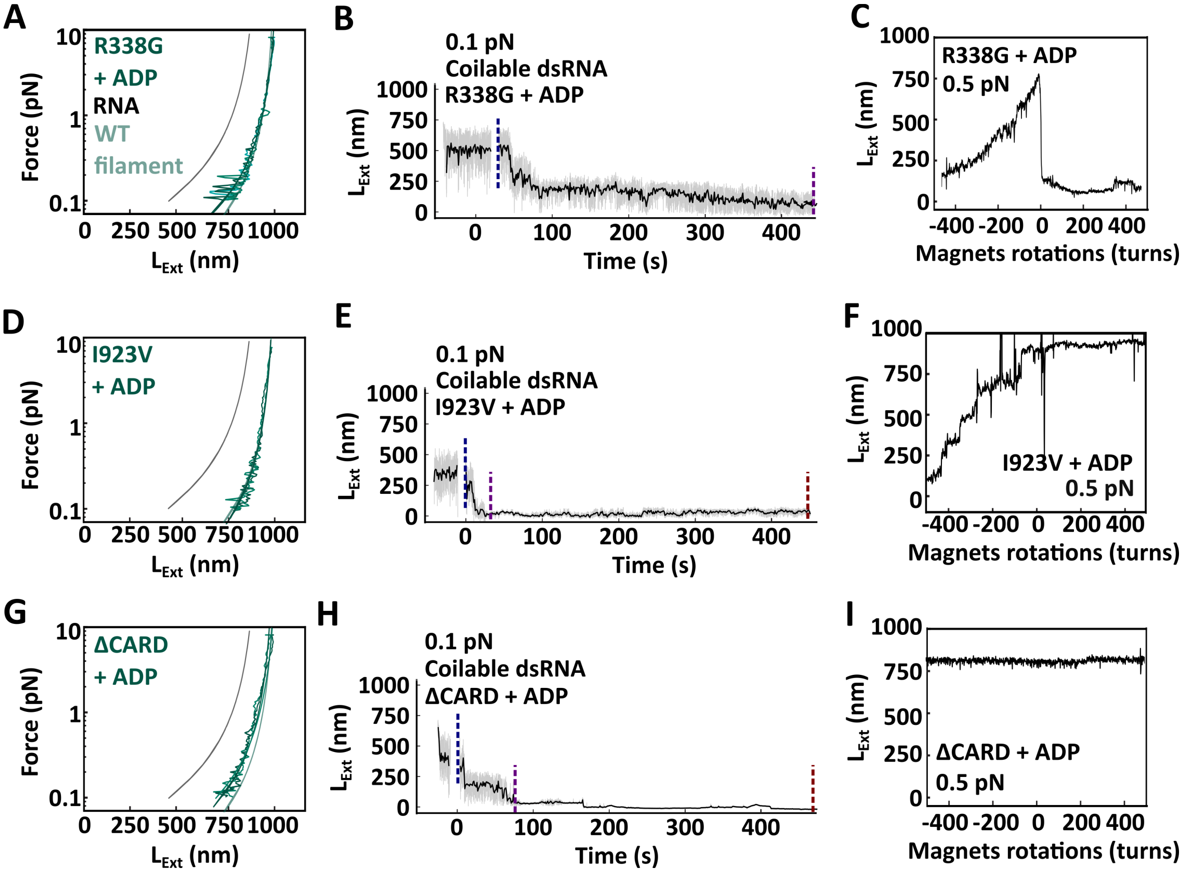


**Figure S5: Filament formation with either MDA5 ΔCARD or MDA5 R338G is similar to WT MDA5. (A)** Force-extension traces of MDA5 R338G filaments formed with ADP (green). **(B)** Time trace filament formation on a coilable dsRNA tether with 100 nM MDA5 R338G and 2 mM ADP. **(C)** Rotation-extension trace of MDA5 R338G filament assembled with ADP**. (D)** Force-extension traces of MDA5 I923V filaments formed with ADP (green). **(E)** Time trace of filament formation on a coilable dsRNA tether with 100 nM MDA5 I923V and 2 mM ADP. **(F)** Rotation-extension of MDA5 I923V filament. **(G)** Force-extension traces of MDA5 ΔCARD filaments formed with ADP (green). **(H)** Time trace of filament assembly in the presence of ADP on a coilable dsRNA tether with 100 nM MDA5 ΔCARD and 2 mM ADP. **(I)** Rotation-extension trace of MDA5 ΔCARD filament assembled with ADP**.** (A, D, G) The mean force-extension traces of dsRNA (grey line) and WT MDA5 filament (teal line) are represented for comparison. Statistics and non-extensible WLC fit parameters are provided in **Table S7**. (B, E, H) The raw (58 Hz) and time averaged (1 Hz) traces are represented in grey and black, respectively.

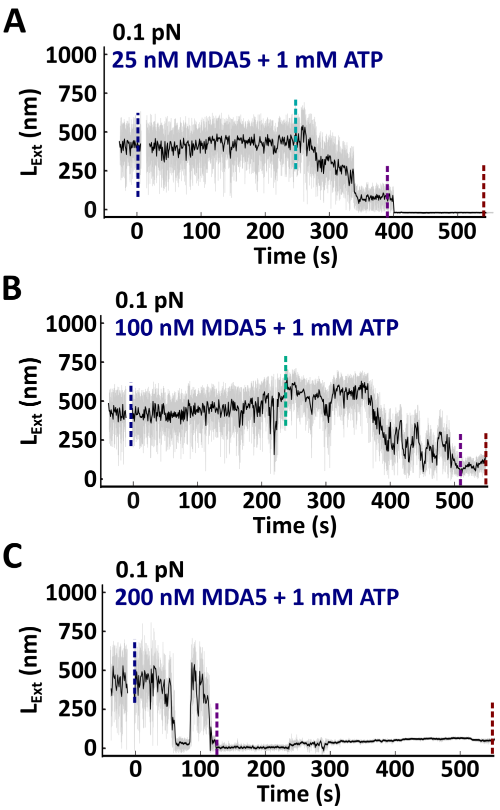


**Figure S6: Assay resolution and filament nucleation, formation and compaction rates as a function of MDA5 concentration in presence of ATP. (A, B, C)** Time traces of MDA5 filament formation in presence of 1 mM ATP and 0.1 pN with **(A)** 25 nM, **(B)** 100 nM, or **(C)** 200 nM MDA5, respectively. The raw (58 Hz) and time averaged (1 Hz) traces are represented in grey and black, respectively. The vertical dashed lines indicate the end of each phase: flushing (dark blue), pre-compaction (teal), compaction (purple) and the compacted filament (red). The raw (58 Hz) and time averaged (1 Hz) traces are represented in grey and black, respectively.

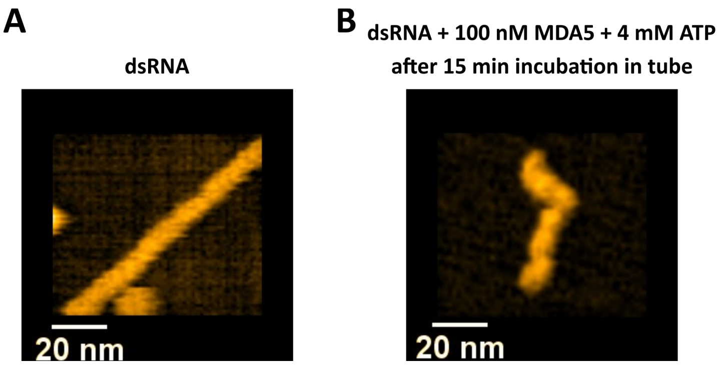


**Figure S7: High-speed AFM images of bare dsRNA and ssRNA. (A)** AFM images taken for a bare dsRNA on mica. **(B)** AFM image of a sample of 100 nM MDA5 with 4 mM ATP and dsRNA after incubation for 15 min in a reaction tube. Only ssRNA strands remained present in the sample. This confirms that in the absence of obstacles blocking the ends of the dsRNA tether, MDA5 can unwind the complete dsRNA. The image reveals only ssRNA remains.

**Video S1:** HS-AFM movie capturing the MDA5 –dsRNA interaction upon 6 mM ATP hydrolysis in real-time. The imaging rate is 300 milliseconds per frame.

**Video S2:** This is an example HS-AFM movie capturing the ssRNA-like coiled structure on mica extracted from a mixture of dsRNA, 100 nM MDA5, and 4 mM ATP. The imaging rate is 500 milliseconds per frame.

**Table S1: Mean filament nucleation and formation rate of MDA5-dsRNA filament in presence of 2 mM AMP-PNP at 0.1 pN**

| [MDA5] (nM) | $k_{nucl}$ | ± | std (1/s) | $k_{form}$ | ± | std (monomers/s) | $k_{form}$ | ± | std (nm/s) | N |
|---|---|---|---|---|---|---|---|---|---|---|
| 25 | 0.010 | ± | 0.001 | 0.7 | ± | 0.1 | 0.8 | ± | 0.1 | 44 |
| 50 | 0.022 | ± | 0.003 | 1.2 | ± | 0.1 | 1.4 | ± | 0.1 | 34 |
| 60 | 0.015 | ± | 0.001 | 1.1 | ± | 0.1 | 1.2 | ± | 0.1 | 98 |
| 75 | 0.019 | ± | 0.002 | 1.4 | ± | 0.1 | 1.6 | ± | 0.1 | 55 |
| 100 | 0.024 | ± | 0.002 | 1.2 | ± | 0.1 | 1.5 | ± | 0.1 | 70 |
| 150 | 0.042 | ± | 0.007 | 3.9 | ± | 0.3 | 4.3 | ± | 0.3 | 53 |
| 200 | 0.042 | ± | 0.004 | 4.5 | ± | 0.4 | 5.0 | ± | 0.4 | 42 |
| 300 | 0.062 | ± | 0.009 | 7.9 | ± | 1.0 | 8.5 | ± | 1.1 | 33 |

**Table S2: Mean contour length and persistence length of MDA5-dsRNA filaments in presence of 2 mM AMP-PNP**

| [MDA5] (nM) | $L_C$ | ± | std (nm) | $L_p$ | ± | std (nm) | N WLC |
|---|---|---|---|---|---|---|---|
| 0 | 888 | ± | 89 | 41 | ± | 11 | 321 |
| 25 | 938 | ± | 66 | 90 | ± | 28 | 58 |
| 50 | 929 | ± | 89 | 105 | ± | 24 | 135 |
| 60 | 816 | ± | 42 | 108 | ± | 28 | 144 |
| 75 | 908 | ± | 85 | 161 | ± | 55 | 103 |
| 100 | 848 | ± | 68 | 205 | ± | 33 | 39 |
| 150 | 826 | ± | 100 | 216 | ± | 59 | 19 |
| 200 | 853 | ± | 78 | 194 | ± | 47 | 14 |
| 300 | 972 | ± | 73 | 132 | ± | 63 | 40 |

**Table S3: Contour length and persistence length of MDA5-dsRNA filament after 30 min intervals**

| | 30 min | | 60 min | | 90 min | | 120 min | | N |
|---|---|---|---|---|---|---|---|---|---|
| | $L_C$ ± std (nm) | $L_P$ ± std (nm) | $L_C$ ± std (nm) | $L_P$ ± std (nm) | $L_C$ ± std (nm) | $L_P$ ± std (nm) | $L_C$ ± std (nm) | $L_P$ ± std (nm) | |
| 2 mM AMP-PNP | 929 ± 137 | 153 ± 87 | 914 ± 115 | 193 ± 93 | 911 ± 109 | 203 ± 93 | 916 ± 127 | 191 ± 81 | 16 |
| 2 mM ADP | 998 ± 57 | 98 ± 92 | 961 ± 73 | 122 ± 11 | 1002 ± 56 | 171 ± 103 | 940 ± 73 | 168 ± 60 | 5 |
| No nucleotide | 965 ± 62 | 198 ± 147 | 939 ± 41 | 227 ± 86 | 933 ± 37 | 153 ± 42 | 933 ± 43 | 245 ± 143 | 2 |
| 700 mM NaCl | 908 ± 94 | 56 ± 19 | - | - | - | - | - | - | 17 |

**Table S4: Rotation rates of MDA5 WT-dsRNA filaments upon negative and positive supercoiling**

| Force (pN) | Negative $k_{rot}$ | ± | std (nm/turn) | N | Positive $k_{rot}$ | ± | std (nm/turn) | N |
|---|---|---|---|---|---|---|---|---|
| 0.5 | 1.49 | ± | 0.15 | 15 | 14.96 | ± | 1.83 | 13 |
| 1 | 1.20 | ± | 0.09 | 18 | 6.32 | ± | 0.95 | 16 |
| 2 | 1.04 | ± | 0.03 | 24 | 2.07 | ± | 0.56 | 22 |
| 4 | 0.94 | ± | 0.02 | 10 | 0.59 | ± | 0.11 | 8 |
| 8 | 0.71 | ± | 0.03 | 24 | 0.74 | ± | 0.06 | 22 |

**Table S5: Mean filament formation and compaction rate of MDA5-dsRNA filament in presence of 2 mM AMP-PNP at 0.1 pN on a coilable dsRNA tether.**

| [MDA5] (nM) | $k_{form}$ | ± | std (nm/s) | $k_{comp}$ | ± | std (nm/s) | | N |
|---|---|---|---|---|---|---|---|---|
| 100 | 0.1 | ± | 0.3 | 5.2 | ± | 0.9 | | 6 |

**Table S6: Compaction rates and reversal counts of MDA5 WT-dsRNA filaments**

| [ATP] (mM) | Force (pN) | $k_{comp}$ | ± | std (nm/s) | rel. count reversal events | ± | std | N |
|---|---|---|---|---|---|---|---|---|
| 1 | 0.1 | 1.35 | ± | 0.19 | | | | 67 |
| 0.5 | 0.5 | 0.40 | ± | 0.02 | 0.4 | ± | 0.8 | 102 |
| 1 | | 0.48 | ± | 0.03 | 0.7 | ± | 1.5 | 77 |
| 2 | | 0.72 | ± | 0.04 | 2.7 | ± | 2.9 | 55 |
| 3 | | 0.95 | ± | 0.05 | 2.2 | ± | 2.3 | 52 |
| 4 | | 1.53 | ± | 0.08 | 1.8 | ± | 2.2 | 95 |
| 0.5 | 1 | 0.22 | ± | 0.01 | 0.4 | ± | 1.0 | 44 |
| 1 | | 0.32 | ± | 0.03 | 0.3 | ± | 1.1 | 33 |
| 2 | | 0.68 | ± | 0.03 | 2.4 | ± | 2.7 | 59 |
| 3 | | 0.81 | ± | 0.03 | 2.0 | ± | 2.3 | 81 |
| 4 | | 1.29 | ± | 0.06 | 1.7 | ± | 2.3 | 64 |
| 0.5 | 2 | 0.20 | ± | 0.01 | 0.0 | ± | 0.1 | 46 |
| 1 | | 0.23 | ± | 0.01 | 0.2 | ± | 0.8 | 91 |
| 2 | | 0.50 | ± | 0.02 | 0.4 | ± | 0.8 | 41 |
| 3 | | 0.76 | ± | 0.03 | 0.3 | ± | 0.5 | 68 |
| 4 | | 0.52 | ± | 0.02 | 0.2 | ± | 0.5 | 84 |
| 0.5 | 4 | 0.15 | ± | 0.01 | 0.1 | ± | 0.5 | 57 |
| 1 | | 0.19 | ± | 0.01 | 0.5 | ± | 0.7 | 43 |
| 2 | | 0.29 | ± | 0.01 | 0.2 | ± | 0.6 | 62 |
| 3 | | 0.41 | ± | 0.01 | 0.3 | ± | 0.7 | 51 |
| 4 | | 0.58 | ± | 0.01 | 0.4 | ± | 1.1 | 163 |

**Table S7: Mean persistence length and mean contour length of MDA5 mutant-dsRNA filaments**

| | $L_C \pm$ std (nm) | $L_P \pm$ std (nm) | N | $L_C \pm$ std (nm) | $L_P \pm$ std (nm) | N |
|---|---|---|---|---|---|---|
| | 2 mM ADP | | | 4 mM ATP | | |
| R338G | 886 ± 18 | 190 ± 18 | 66 | 865 ± 23 | 190 ± 20 | 66 |
| I923V | 885 ± 9 | 150 ± 23 | 177 | - | - | |
| ΔCARD | 945 ± 23 | 185 ± 19 | 68 | 886 ± 18 | 42 ± 2 | 68 |

**Table S8: Trace classification of 50 nM MDA5 on gapped RNA in presence of 1 mM ATP at 1pN**

| RNA constructs | Extension | | | Dissociation | | | Partial Compaction | | | Full Compaction | | | Total |
|---|---|---|---|---|---|---|---|---|---|---|---|---|---|
| | Mean | ± | SEM | Mean | ± | SEM | Mean | ± | SEM | Mean | ± | SEM | N |
| dsRNA | 5.6% | ± | 7.5% | 0.0% | ± | 0.0% | 27.8% | ± | 14.6% | 66.7% | ± | 15.4% | 36 |
| 3'-gap up | 20.8% | ± | 16.2% | 8.3% | ± | 11.1% | 25.0% | ± | 17.3% | 45.8% | ± | 19.9% | 24 |
| 3'-gap down | 31.0% | ± | 11.9% | 3.4% | ± | 4.7% | 36.2% | ± | 12.4% | 29.3% | ± | 11.7% | 58 |
| 3'-gap both | 56.8% | ± | 16.0% | 8.1% | ± | 8.8% | 32.4% | ± | 15.1% | 2.7% | ± | 5.2% | 37 |
| 5'-gap down | 25.0% | ± | 16.0% | 7.1% | ± | 9.5% | 21.4% | ± | 15.2% | 46.4% | ± | 18.5% | 28 |
| 5'-gap up | 30.8% | ± | 10.2% | 1.3% | ± | 2.5% | 39.7% | ± | 10.9% | 28.2% | ± | 10.0% | 78 |
| 5'-gap both | 8.5% | ± | 6.0% | 28.0% | ± | 9.7% | 19.5% | ± | 8.6% | 43.9% | ± | 10.7% | 82 |

**Table S9: Trace classification of 400 nM MDA5 on gapped RNA in presence of 1 mM ATP at 1pN**

| RNA constructs | Extension | | | Dissociation | | | Compaction | | | Full Compaction | | | Total |
|---|---|---|---|---|---|---|---|---|---|---|---|---|---|
| | Mean | ± | SEM | Mean | ± | SEM | Mean | ± | SEM | Mean | ± | SEM | N |
| dsRNA | 8.3% | ± | 11.1% | 8.3% | ± | 11.1% | 16.7% | ± | 14.9% | 66.7% | ± | 18.9% | 24 |
| 3'-gap up | 21.4% | ± | 15.2% | 3.6% | ± | 6.9% | 57.1% | ± | 18.3% | 17.9% | ± | 14.2% | 28 |
| 3'-gap down | 14.9% | ± | 8.1% | 5.4% | ± | 5.2% | 50.0% | ± | 11.4% | 29.7% | ± | 10.4% | 74 |
| 3'-gap both | 9.5% | ± | 12.6% | 0.0% | ± | 0.0% | 57.1% | ± | 21.2% | 33.3% | ± | 20.2% | 21 |
| 5'-gap down | 60.0% | ± | 42.9% | 0.0% | ± | 0.0% | 20.0% | ± | 35.1% | 20.0% | ± | 35.1% | 5 |
| 5'-gap up | 48.7% | ± | 15.7% | 2.6% | ± | 5.0% | 38.5% | ± | 15.3% | 10.3% | ± | 9.5% | 39 |
| 5'-gap both | 29.4% | ± | 21.7% | 0.0% | ± | 0.0% | 41.2% | ± | 23.4% | 29.4% | ± | 21.7% | 17 |

**Table S10: Count of coilable and non-coilable tethers before and after RNase A treatment**

| | coilable before | Still coilable after | Lost bead after | Non-coilable after |
|---|---|---|---|---|
| With RNase A | 31 | 0 | 6 | 25 |
| Without ATP | 16 | 12 | 4 | 0 |
| Without RNase A | 21 | 13 | 7 | 1 |

**Table S11: Primers for RNA constructs**

| | |
|---|---|
| 5'-3' strand BIO handle fw | GGATCCTACCTGACGCTTTT |
| 5'-3' strand BIO handle rev | TAATACGACTCACTATAGGCAAACGGCTTGATATCC |
| 5'-3' strand strand fw | ATTCAGGGACTGCCGATG |
| 5'-3' strand strand rev | TAATACGACTCACTATAGGACGTTTCGGATCTTCC |
| 5'-3' strand DIG handle fw | GGAGCGTAAAATTCAGTTCTTC |
| 5'-3' strand DIG handle rev | TAATACGACTCACTATAGGTTAACCTCAACTTCCATTTCC |
| 5'-3' strand 3'-gap up fw | TCGTGTCACCCAGTCGGACC |
| 5'-3' strand 3'-gap up rev | TAATACGACTCACTATAGGAGCGCCGCTTCCATGTCCTGGAACGCT |
| 5'-3' strand 5'-gap down fw | TAATACGACTCACTATAGGCACGTCGCGCAGATG |
| 3'-5' strand 3'-gap down fw | TAATACGACTCACTATAGGCGGATAACGAACATCTG |
| 3'-5' strand 3'-gap down rev | TCCGACATGGAACGCAA |
| 3'-5' strand rev | GGTTAACCTCAACTTCCATTTCC |
| 3'-5' strand fw | TAATACGACTCACTATAGGATCCTACCTGACGCTTTT |
| 3'-5' strand 5'-gap up fw | TAATACGACTCACTATAGGCGCGCGTGCTGC |
| 3'-5' strand 5'-gap up rev | TGTGTCGGCTGCACCGAC |
| 3'-5' strand DIG handle for 3'-gap up fw | TAATACGACTCACTATAGGTCGGATAAGGCGTTAGG |
| 3'-5' strand 3'-gap up rev | ATGGAACGCAAATCATCAGC |
